# Stepwise evolution of *E. coli* C and ΦX174 reveals unexpected lipopolysaccharide (LPS) diversity

**DOI:** 10.1101/2022.09.06.506728

**Authors:** Jordan Romeyer Dherbey, Lavisha Parab, Jenna Gallie, Frederic Bertels

## Abstract

Phage therapy is a promising method for the treatment of multi-drug-resistant bacterial infections. However, its long-term efficacy depends on understanding the evolutionary effects of the treatment. Current knowledge of such evolutionary effects is lacking, even in well-studied systems. We used the bacterium *Escherichia coli* C and its bacteriophage ΦX174, which infects cells using host lipopolysaccharide (LPS) molecules. We first generated 31 bacterial mutants resistant to ΦX174 infection. Based on the genes disrupted by these mutations, we predicted that these *E. coli* C mutants collectively produce eight unique LPS structures. We then developed a series of evolution experiments to select for ΦX174 mutants capable of infecting the resistant strains. During phage adaptation, we distinguished two types of phage resistance: one that was easily overcome by ΦX174 with few mutational steps (“easy” resistance), and one that was more difficult to overcome (“hard” resistance). We found that increasing the diversity of the host and phage populations could accelerate the adaptation of phage ΦX174 to overcome the hard resistance phenotype. From these experiments, we isolated 16 ΦX174 mutants that, together, can infect all 31 initially resistant *E. coli* C mutants. Upon determining the infectivity profiles of these 16 evolved phages, we uncovered 14 distinct profiles. Given that only eight profiles are anticipated if the LPS predictions are correct, our findings highlight that the current understanding of LPS biology is insufficient to accurately forecast the evolutionary outcomes of bacterial populations infected by phage.

## Introduction

Multidrug-resistant bacterial infections are one of the most pressing issues in medicine, a situation that is only expected to worsen in the coming decades (Centers for Disease Control and Prevention (U.S.) 2019; WHO 2021; Murray et al. 2022). An alternative to treating bacterial infections with antibiotics is phage therapy. Major research efforts are being conducted to develop efficient therapies (Oechslin 2018; Bull et al. 2019; Monteiro et al. 2019). An efficient therapy should be able to eliminate a pathogen without selecting for resistance to the phage being used to treat that pathogen. Phage resistance can often evolve by modifying the cell envelope to avoid phage adsorption. The outer membrane of all Gram-negative bacteria is covered in lipo<u>p</u>oly<u>s</u>accharides (LPSs), which is consequently the first contact point between a bacterium and its environment (Zhang et al. 2013). LPS structure determines whether a bacterium will be recognized by the immune system, the efficiency of nutrient uptake as well as the susceptibility to antibiotics and, importantly, the susceptibility to phage infection (Nikaido 2003; Simpson and Trent 2019; Mutalik et al. 2020). Hence, LPS structure is an important factor determining the success of phage therapy.

The functions of genes involved in synthesizing and assembling bacterial LPS have been elucidated (Schnaitman and Klena 1993; Amor et al. 2000; Raetz and Whitfield 2002; Qian et al. 2014; Klein and Raina 2019). Most *Escherichia coli* strains produce LPS molecules composed of three parts: the lipid A, the core <u>o</u>ligo<u>s</u>accharide (OS), and the O-antigen chain. The LPS core OS itself consists of a very conserved inner core across *E. coli* species and a variable outer core (Amor et al. 2000). Strains with the tripartite LPS molecules (lipid A, core OS, and O-antigen) are termed “smooth” (Raetz and Whitfield 2002). Some *E. coli* strains, however, produce LPS molecules that lack one or more components. LPS molecules lacking the O-antigen are classified as “rough”, and molecules lacking both the O-antigen and the outer core OS as “deep rough” (van der Ley et al. 1986; Schnaitman and Klena 1993; Yethon et al. 1998; Whitfield et al. 1999; Amor et al. 2000; Klein et al. 2013). These modifications in LPS structure can lead to a plethora of phenotypic effects (Simpson and Trent 2019), including (i) destabilization of the outer membrane (Raetz and Whitfield 2002), (ii) changes in the expression of some outer membrane proteins, (iii) modification of intracellular turgor pressure (Pagnout et al. 2019), (iv) increased susceptibility to hydrophobic compounds such as antimicrobial peptides, antibiotics, and bacteriocins (van der Ley et al. 1986; Schnaitman and Klena 1993; Yethon et al. 1998; Whitfield et al. 1999; Amor et al. 2000; Klein et al. 2013), (v) altered interactions with the host immune system (Raetz and Whitfield 2002; Matsuura 2013; Maldonado et al. 2016), (vi) alteration of the redox status of cells leading to oxidative stress (Seregina et al. 2022), and (vii) changes in resistance to phages (Hancock and Reeves 1976; Labrie et al. 2010; Kulikov et al. 2019; Mutalik et al. 2020).

Phage ΦX174 is a small (∼320 Å), tailless, lytic coliphage from the *Microviridae* family. It carries a circular, single-stranded DNA genome of only 5386 bases long that encodes 11 genes (Sinsheimer 1959; Sanger et al. 1978). To enter a host cell, ΦX174 relies solely on attaching to the core OS by one of its spike proteins before migrating and irreversibly adsorbing to the host’s cell surface via its capsid (Incardona and Selvidge 1973; Incardona et al. 1985; Sun et al. 2017). ΦX174 does it with a high degree of specificity. For instance, among 783 different *E. coli* isolates, only six (0.8 %) could be infected by wildtype ΦX174 (Michel et al. 2010). In the laboratory, ΦX174 infects - and hence is usually grown on - *E. coli* C, which produces rough type (*i.e.*, lacking in O-antigen) LPS molecules (Feige, Ulrich and Stirm, Stephan 1976; Inagaki et al. 2000; Kawaura et al. 2000; Król et al. 2019) (**Figs. 1 and S1**).

**Fig. 1.**
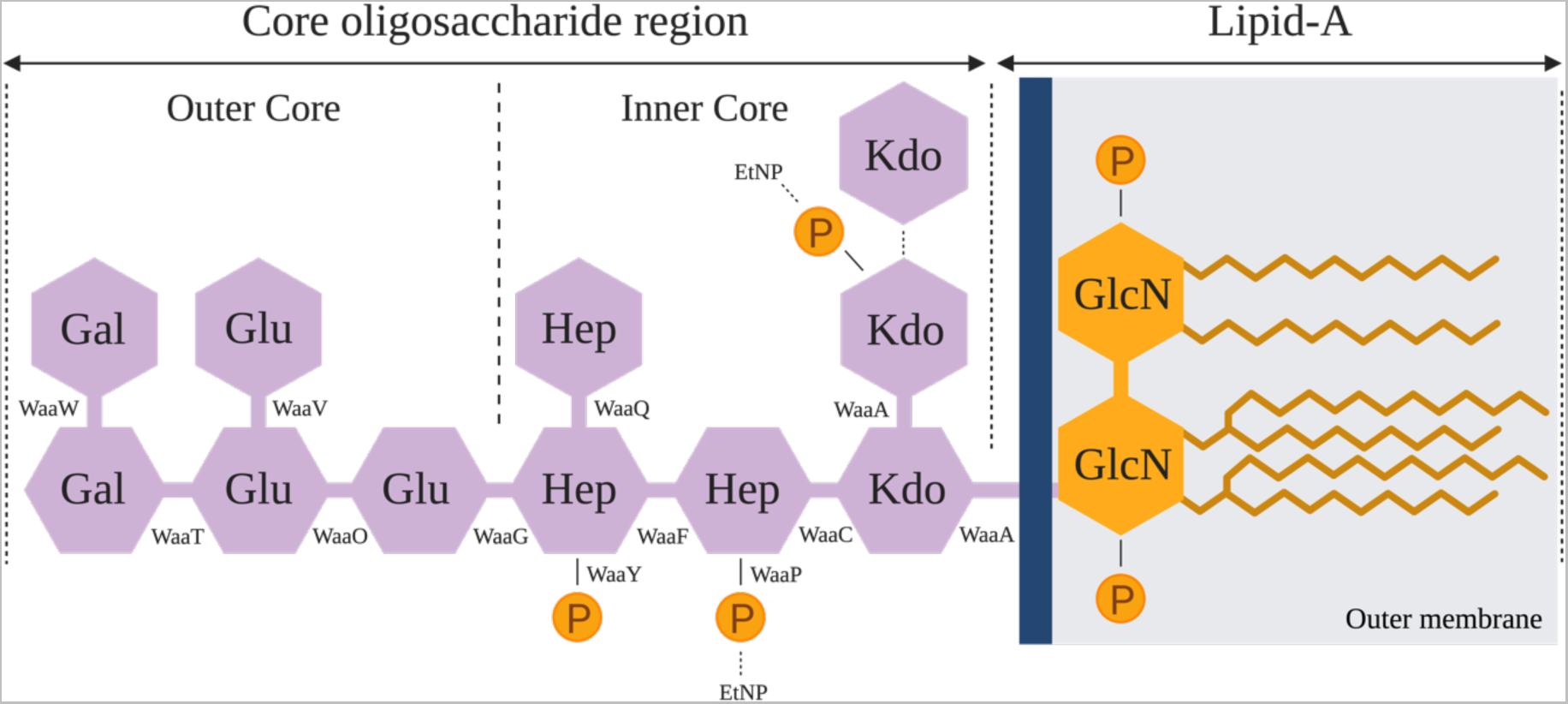
Structure of *E. coli* C’s rough type LPS. The rough type LPS of *E. coli* C is composed of two parts: lipid A (composed of an acetylated and 1,4’-diphosporylated β(1→6)-linked glucosamine (GlcN) disaccharide), and the core oligosaccharide (Raetz and Whitfield 2002). The core oligosaccharide is subdivided into a structurally conserved inner core and an outer core. The dashed lines show non-stochiometric substitutions of phosphate (P), ethanolamine (EtNP), and 3-deoxy-D-*manno*-octulosonic acid (Kdo) residues on the LPS (Kojima et al. 2010). Details of the LPS assembly process are provided in **Fig. S1**.

Here, we explore the evolutionary potential of *E. coli* C to become resistant to infection by ΦX174 and, conversely, the potential of ΦX174 mutants to infect phage-resistant *E. coli* C hosts with modified LPS structures. We first generated a set of 31 *E. coli* C mutants that are resistant to infection by ΦX174 wildtype and determined that each mutant carries at least one mutation affecting LPS biosynthesis.

The currently accepted model for predicting LPS structure posits that the presence or absence of each LPS gene leads to a single LPS phenotype. Based on the position of each mutation, the one-gene-one-phenotype model allows us to predict the LPS structure of each of our LPS mutants. Our 31 *E. coli* C mutants collectively produce LPS molecules with eight distinct structures. To test the accuracy of these predictions, we evolved and genotypically characterized a set of 16 ΦX174 genotypes that can, together, infect all initially resistant 31 *E. coli* C mutants. If the LPS structure predictions are correct, then all bacteria predicted to carry the same LPS structure should be susceptible to the same set of phages. Interestingly, this was not always the case, indicating that the diversity of LPS structures generated by LPS-pathway mutations is much higher than anticipated from previous studies. Bacterial strains with different mutations in the same LPS gene sometimes have strikingly different infection phenotypes. Even the required number of phage mutations to overcome these resistances can differ significantly. A better understanding of the evolution of phage resistance hence requires a better understanding of the biology and evolution of LPS structures.

## Results

### Diverse LPS structures confer resistance to ΦX174 infection

To study LPS diversity in *E. coli* C, we generated 35 spontaneous phage-resistant strains and used whole genome re-sequencing to identify the mutations conferring resistance (see **Methods**). Due to a lack of isogeneity or only partial resistance to wildtype ΦX174, four of these strains (*E. coli* C R1, R3, R15, and R19) were excluded from downstream analyses (see **Text S1**). Of the remaining 31 strains, 27 are predicted to carry a single mutation, three carry two mutations, and one carries three mutations (see **Table S1**). A total of 36 mutations were identified, 32 of which are unique (including 15 nucleotide substitutions, 13 deletions, one duplication, and three IS*4* and IS*5* insertion events). Twenty-four (of 32, or 75%) mutations introduce premature stop codons or lead to frameshifts and thus are highly likely to disrupt gene function.

Interestingly, two mutations are shared by two strains (R27/R31 and R8/R10), and one mutation is shared by three strains (R4/R24/R29). Given that there are almost limitless ways to disrupt gene function, this high degree of parallelism is initially surprising. We note that the repetition of these mutations is unlikely to result from their positive selection. That is, phage-resistant *E. coli* C mutants of both low and high fitness are expected to have an equal chance of appearing on our agar plates as long as they can form visible colonies (Luria and Delbrück 1943). Presumably, the observed parallelism is, instead, due to elevated mutation rates at particular genomic positions (Lind et al. 2019). The fact that two of three parallel mutations occur in homopolymeric tracts supports this hypothesis (Moxon et al. 1994).

Mutations in genes encoding the LPS machinery are expected to lead to changes in LPS structure and, in some cases, a switch from a rough to a deep rough phenotype (Yethon et al. 1998; Whitfield et al. 1999; Yethon et al. 2000). It has been shown that logical predictions of mutant LPS structures can be made. If an enzyme that synthesizes a particular LPS component is absent or non-functional, then that LPS component cannot be synthesized. In addition, LPS components assembled downstream of the missing component are also absent from the final LPS structure. The assumption that each mutation in an LPS gene leads to a complete lack of function allows us to predict the LPS structure and type of each of the 31 *E. coli* C phage-resistant mutants (**Figs. 2**, **S1**, **S2 and Table S1**). A total of seven emergent LPS structures were predicted from our set of mutants (**Fig. 2**). Compared with the *E. coli* C wildtype LPS, three of these seven LPS structures carry alterations in the outer-core LPS and hence are predicted to be of the rough type (as is wildtype *E. coli* C; ten mutants). The remaining four LPS structures differ from wildtype LPS in both the inner- and outer-core LPS, and are presumed to be of the deep rough type (twenty mutants) (see **Table S1**).

**Fig. 2.**
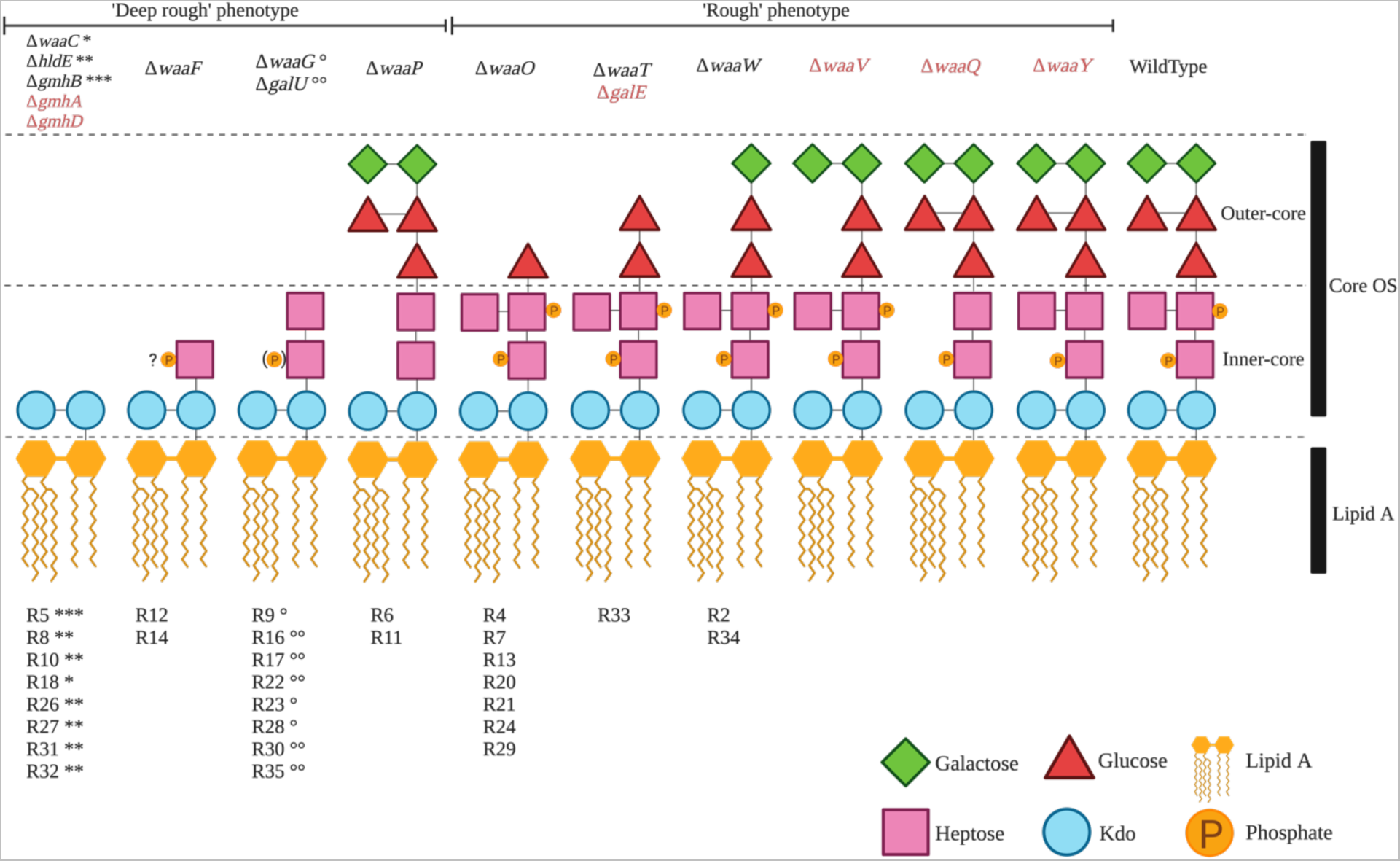
Predicted LPS structures of 30 ΦX174-resistant *E. coli* C strains. Predicted LPS structures fall into two groups depending on their degree of truncation and/or loss of phosphate groups: the “deep rough” phenotype with a completely truncated outer core or a loss of phosphate groups (twenty *E. coli* C mutants) and the “rough” phenotype with smaller LPS truncations located in the outer core (ten *E. coli* C mutants). The LPS phenotype of the final *E. coli* C mutant (R25) could not be predicted (see text). The products of genes in red are also involved in LPS biosynthesis and, although no mutations were identified in these genes during this study, are also potential targets for phage resistance. Details on the functions of *gmhA*, *gmhB*, *gmhD*, *hldE*, *galU*, and *galE* are provided in **Fig. S2**. “?P”: No information was found on the phosphorylation of Hep(I) in the absence of *waaF*; (P): only 40% of the hexose phosphorylation was observed (Yethon et al. 2000); R#: the number gave to each *E. coli* C resistant strain; “*, **, ***, °, °°”: symbols link the resistant strains (R#) to particular mutated LPS genes (Δ*gene-names*). Resistant strains with identical symbols carry their mutations in the same LPS gene. For example, [R8, R10, R26, R27, R31, and R32]-strains each carry a mutation in the *hldE* gene, and [R9, R23, and R28]-strains all carry a mutation in the *waaG* gene (**Table S1**).

The LPS structure and type of the remaining mutant (*E. coli* C R25) were not predicted because the effect of the phage-resistance mutation on LPS biosynthesis remains unclear. R25 carries a nonsense mutation in *rfaH*. The *rfaH* gene encodes a transcriptional antiterminator that regulates the *waa* operon (Bailey et al. 1997; Klein and Raina 2019), meaning that any number of LPS biosynthetic genes could potentially be affected by this mutation.

The LPS structure predictions above can be tested by evolving wildtype ΦX174 to specifically infect. *E. coli* C cells with modified LPS structures. That is, a phage that can infect a given modified LPS structure is expected to be able to infect all bacterial strains displaying this LPS type, regardless of the underlying mutations. Alternatively, the inability of mutant phages to cross-infect bacterial strains of the same predicted LPS structural class (but via different mutations) would indicate that these mutations actually lead to distinct LPS structures.

### ΦX174 evolves to overcome host resistance, a process that can be accelerated by host diversity

Phages that can infect resistant *E. coli* C mutants were evolved by serially transferring wildtype ΦX174 on each of the 31 resistant *E. coli* C strains (**Fig. 3A and Methods**). To reduce the complexity of the experiment, bacterial cultures were not allowed to co-evolve. Instead, phages were inoculated into fresh, exponentially growing host cultures at each transfer. The experiment gave rise to 12 evolved phages that were collectively able to infect 21 (of the 31) initially resistant *E. coli* C strains (these display the “easy” resistant phenotype; see **Table S1**). However, even after 21 transfers, ten (of 31) resistant *E. coli* C strains remained uninfected (the “hard” resistant phenotype; see **Table S1**).

**Fig. 3.**
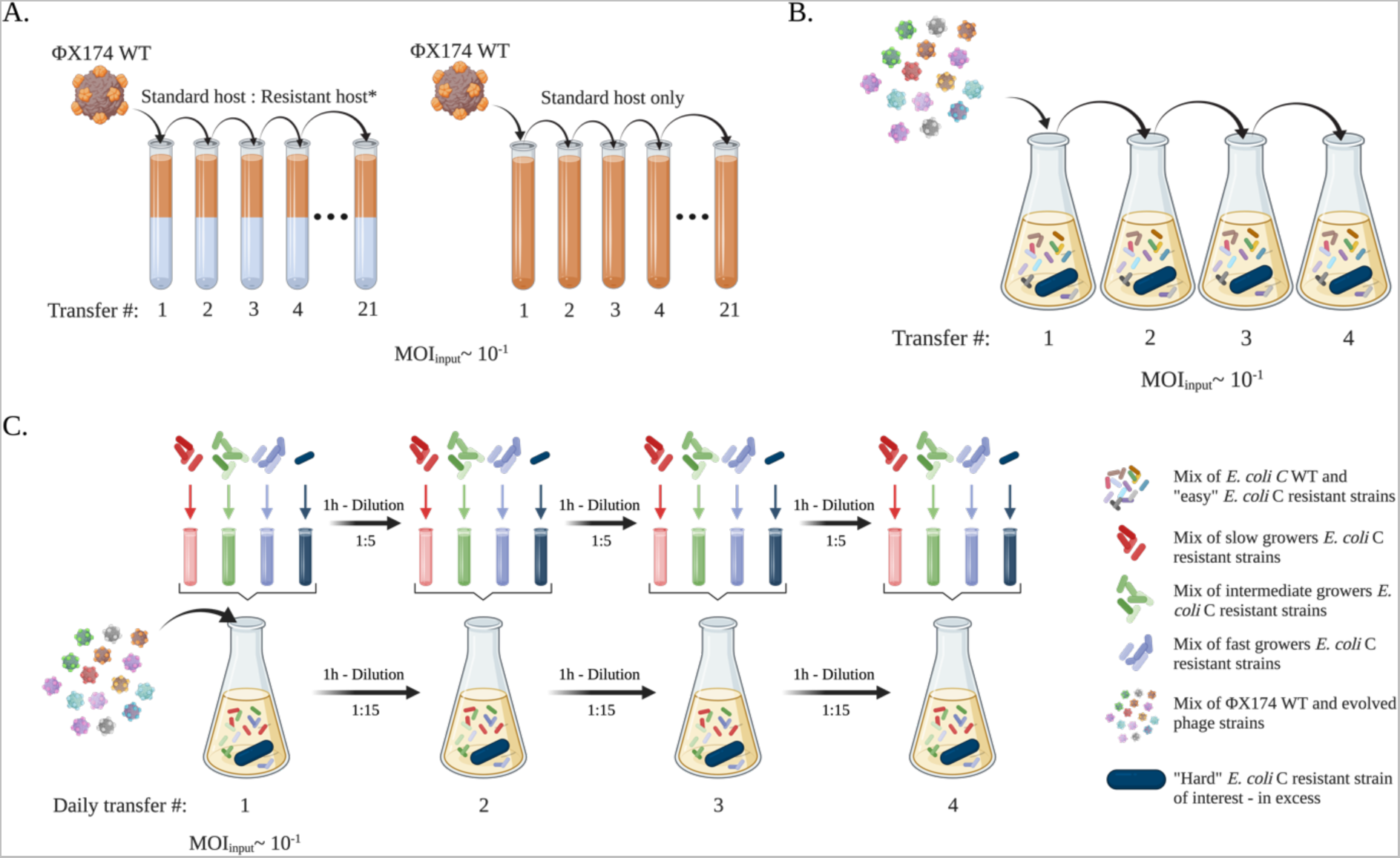
Three phage evolution experiments used to overcome *E. coli* C resistance to wildtype ΦX174. **A.** First evolution experiment (standard): 31 independent lineages, each founded by ΦX174 wildtype, were serially transferred daily for up to 21 days (21 transfers). Phages were grown on non-evolving (i.e., freshly prepared at each transfer) host cultures containing a mixture of *E. coli* C wildtype and one of the 31 initially resistant strains (see **Table S1**). The presence of *E. coli* C wildtype was necessary to propagate ΦX174 and avoid its dilution from one transfer to another. Phages capable of infecting the initially resistant host strain were isolated from 12 lineages. Three control lineages were included, where ΦX174 was cultured on only *E. coli* C wildtype. **B.** Second evolution experiment (increased diversity). A cocktail of ΦX174 wildtype and 14 evolved ΦX174 strains was generated (panel A – the two evolved phages infecting R5 and R19 were part of the final cocktail but removed from the mutational analysis, see **Text S1**). The phage cocktail was transferred daily for up to four days, on non-evolving host cultures containing: (i) *E. coli* C wildtype, (ii) the 14 host strains for which resistance had been overcome (permissive hosts), and (iii) an excess of one of the still-resistant strain of interest (non-permissive host). Phages capable of infecting two resistant strains of interest (*galU* and *waaG* mutants) were isolated from two lines. **C.** Third evolution experiment (increased diversity and generations). The same cocktail used for the second evolution experiment (panel B) was transferred four times daily for four days (i.e., a maximum of 16 transfers) on non-evolving host cultures containing: (i) *E. coli* C wildtype, (ii) the 14 strains for which resistance had been overcome (permissive hosts), and (iii) one of the still-resistant strain of interest (non-permissive host). Notably, all transfers completed on the same day involved transferring both phage and bacteria; supernatants were collected on every fourth transfer, and only phages were transferred. Phages capable of infecting two resistant strains of interest (*waaP/pssA* mutants) were isolated from two lines. Details of the evolved phages are presented in **Table S2**, and further methodological details are provided in **Methods**.

Overcoming phage resistance of this subset of hard-resistant strains may not be achievable with few mutational steps. Hence, either the evolution experiment has to be performed for a much longer period of time, or an experiment that increases the genetic variation available to evolution has to be designed (Hermisson and Pennings 2005; Barrett and Schluter 2008). We designed two further evolution experiments, one to increase genetic diversity and another one to additionally increase the speed of evolution by increasing the number of phage generations per day (**Figs. 3B and 3C**). In both of these experiments, phage diversity was increased by combining all evolved phage strains that successfully reinfected their corresponding resistant strains into a single phage cocktail (Hermisson and Pennings 2005; Barrett and Schluter 2008). The phage cocktail was serially transferred once per day in the overall second evolution experiment, while four transfers per day were performed in the third evolution experiment. Phage diversity was maintained from one transfer to another by adding all corresponding resistant strains to the host culture (see **Methods**). At the end of the evolution experiments, all resistant strains could be cross-infected by at least one of the evolved phage mutants. Our method of evolving phages with an extended infection range is simple and effective and allows the successful evolution and retrieval of phages that, together, can infect all of the 31 resistant *E. coli* C phenotypes.

Increasing both phage and host diversities in the second and third evolution experiments accelerated the adaptation of ΦX174 to resistant *E. coli* C strains. While our experiment quickly gave rise to phages able to infect most non-permissive strains (**Fig. 4A**), some phage-resistant hosts could only be infected when both host and phage diversities were increased (**Figs. 4B** and **4C**).

**Fig. 4.**
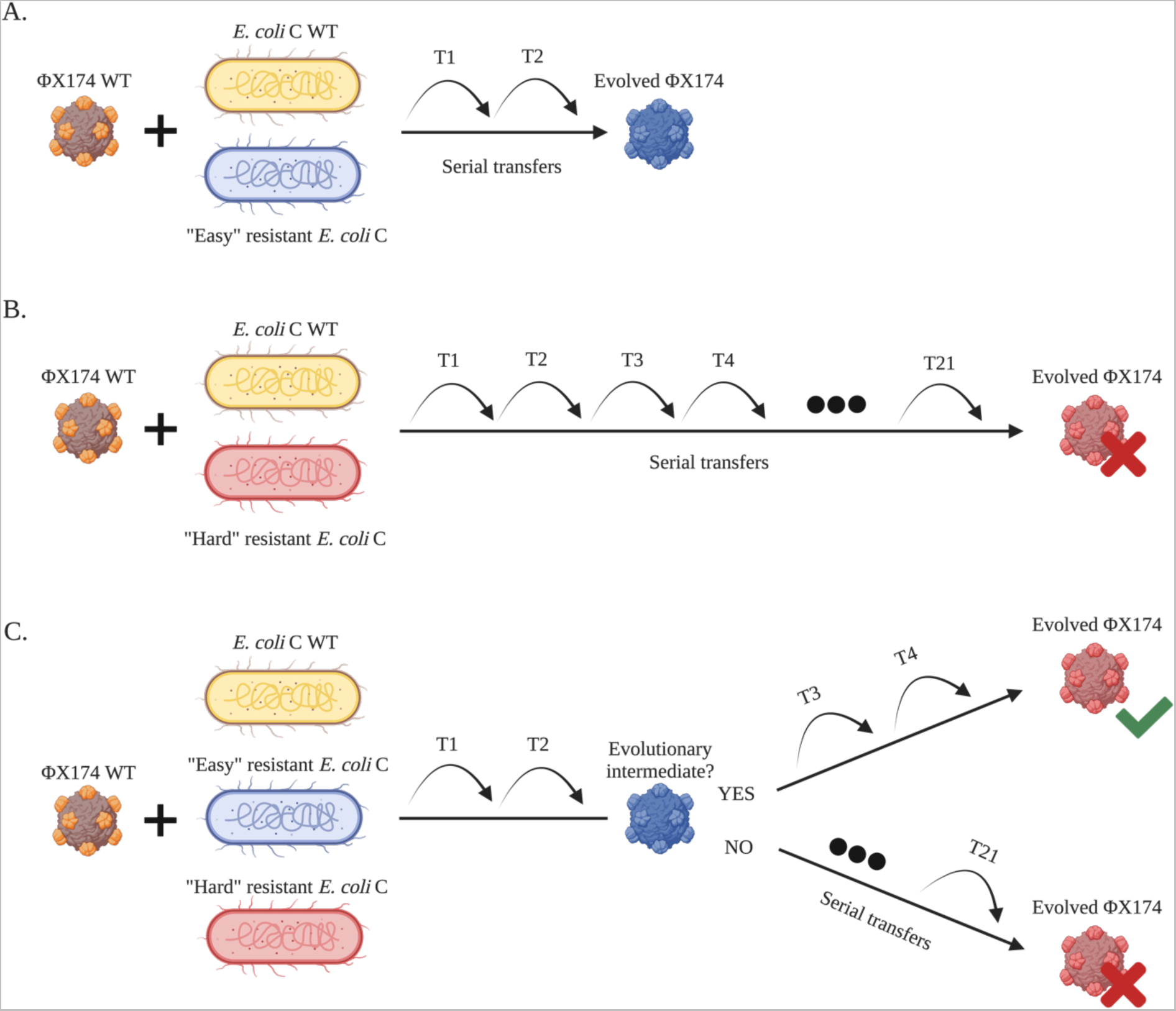
Increased host diversity facilitates viral adaptation. **A.** ΦX174 wildtype evolves on *E. coli* C wildtype and an easy-resistant *E. coli* C strain. Adaptation to the resistant mutant is fast and only requires a few transfers. **B.** ΦX174 wildtype evolves on *E. coli* C wildtype and a hard-resistant *E. coli* C strain that it failed to infect within 21 standard serial transfers. Infecting this resistant strain may be difficult and take a long time. **C.** ΦX174 wildtype evolves in a three-host mix containing *E. coli* C wildtype and two *E. coli* C resistant strains: one easy and one hard. Phage adapts quickly to the easy-resistant phenotype. If the infection of the easy-resistant strain leads to the production of an evolutionary intermediate phage, then the hard phenotype can be infected after a few more transfers. If no evolutionary intermediate phage is produced, then no phage evolves to infect the hard-resistant *E. coli* C strain.

To identify the mutation(s) responsible for the re-infection of the resistant mutants, we performed whole genome re-sequencing on the 16 evolved phages that successfully overcame the corresponding resistance in one of the three evolution experiments (see **Methods**). We found mutations in *F* and/or *H* genes in all 16 evolved phages (**Table S2**). Notably, all hard-resistant *E. coli* C strains - those whose resistance was overcome in the second and/or third evolution experiments - are only infected by phages carrying four mutations (see **Tables S1** and **S2**).

The evolutionary emergence of a multi-mutation phage in a two-strain evolution experiment is expected to require a large number of transfers (**Fig. 4B**). However, this process is significantly accelerated by propagating a diverse mixture of phages. If some of the phages are evolutionary intermediates (i.e., carry a subset of mutations required to overcome hard resistance) then the mutations required to overcome hard resistance can emerge in a single mutational step, either by recombination (Rokyta et al. 2006) or by adding mutation to an existing set (**Fig. 4C**). For example, the hard-resistant strain *E. coli* C R22 (a *galU* mutant; see **Table S1**) is overcome in the second evolution experiment by ΦX174 R22 T1, a phage that carries four mutations (three in *F*, one in *H*; see **Table S2**). Each of these mutations was present in the phage cocktail used to initiate the evolutionary lineage: ΦX174 R18 T4 and ΦX174 R8 T6 contain three of the four mutations, and the final mutation is found in ΦX174 R10 T6. The evolved phage infecting R22 may, therefore, be the product of a recombination event between ΦX174 R10 T6 and ΦX174 R18 T4 (or ΦX174 R8 T6). Alternatively, ΦX174 R18 T4 or ΦX174 R8 T6 could have acquired a single additional mutation. Thus, adaptation to certain easy-resistant LPS phenotypes yields evolutionary intermediate phages that can recombine or acquire a few additional mutations to infect a hard-resistant LPS structure.

Our results show that high host diversity can facilitate the evolution of phages to infect resistant strains. In contrast, Sant et al. showed that high host diversity hinders phage evolution (Sant et al. 2021). The major difference between the experiments is the difficulty of the evolutionary problem posed. Sant et al. only evolved phages to infect easy-resistant strains, strains that could be infected by phages after seven to eight transfers through the acquisition of a few mutations. Evolutionary intermediates will not be required in this situation (**Fig. 4A**). However, as we show here, it is very difficult to infect hard-resistant strains starting with low phage and host diversities (**Fig. 4B**). Hence, increasing the host diversity will accelerate the evolution of phage adaptation under conditions where the additional hosts allow the evolution of intermediate phage phenotypes (**Fig. 4C**).

### ΦX174 overcomes *E. coli* C resistance by mutations in the F capsid and H minor spike proteins

The 16 evolved phages collectively carry 40 mutations, 15 of which are unique (**Fig. 5A and Table S2**). All are non-synonymous and occur in either *F* (encoding the viral capsid) or *H* (encoding the minor spike protein involved in viral DNA injection) genes (**Figs. 5B and 5C**). The majority of phages (13 of 16) carry between two and four mutations (**Fig. 5D**). These results are consistent with previous studies demonstrating that mutations in *F* and *H* are relevant for the adaptation of ΦX174 to novel hosts (Weisbeek et al. 1973; Crill et al. 2000; Cox and Putonti 2010).

**Fig. 5.**
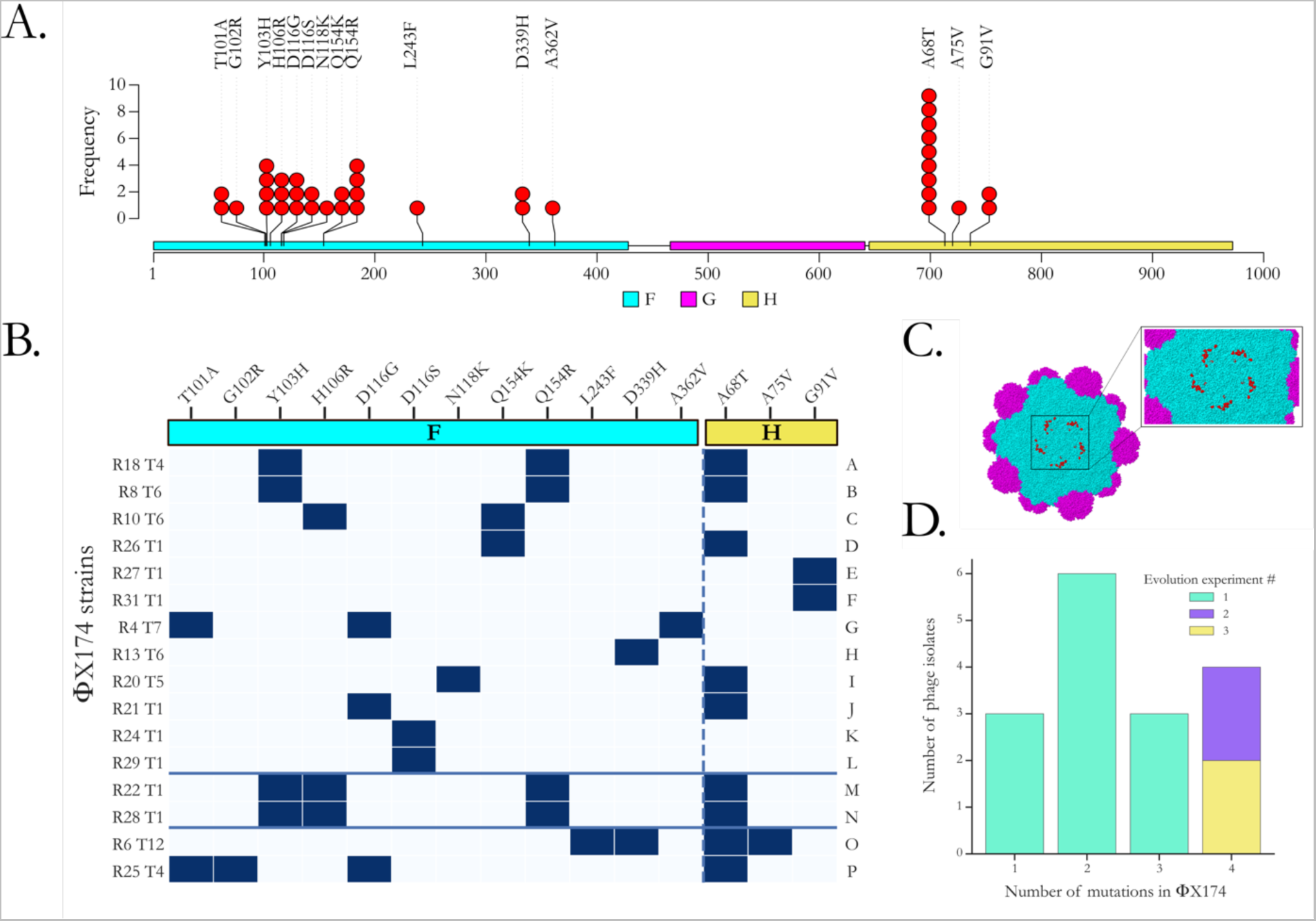
Adaptation of ΦX174 to resistant strains of *E. coli* C. **A.** Positions of all predicted amino acid changes in the F (viral capsid; coloured in cyan) and H (minor spike; coloured in yellow) proteins observed in the 16 evolved ΦX174 strains. No mutation was found in the G protein (major spike; coloured in magenta) despite being an integral part of ΦX174’s recognition system and the first contact point with the LPS structure (Kawaura et al. 2000; Sun et al. 2017). **B.** Substitution matrix of all amino acid changes observed in ΦX174. ΦX174 strains are ordered based on the predicted LPS structures they infected during the evolution experiments (A-F: overcame heptoseless *waaC* and *hldE* mutants; G-L: overcame *waaO* mutants; M-N: overcame *galU* and *waaG* mutants; O: overcame *waaP*/*pssA* mutant; P: overcame *rfaH* mutant). The solid blue lines separate the evolved phages according to the evolution experiment from which they were isolated (A-L: first; M-N: second; O-P: third). The x-axis shows amino acid substitutions in the F and H proteins (separated by the dashed line), R# indicates the resistant bacterial strain the phage evolved on, and T# is the number of transfers required to overcome host resistance. **C.** Location of protein changes on the ΦX174 three-dimensional structure. The F capsid proteins are coloured in cyan, and the G major spike proteins are in magenta. Residues affected by the mutations in this study are in red and shown for one F capsid unit only. The G protein spike (above that pictured here) has been removed to improve visualisation. The image was generated using Geneious Prime (version 2020.1.2) and Protein Data Bank accession code 2BPA (McKenna et al. 1992). Changes in the H protein cannot be displayed as they all occur outside of the currently determined crystallographic structure. **D.** Barplot of the number of evolved phage isolates carrying 1, 2, 3, 4 mutations. Most phage isolates – particularly those isolated from the second and third evolution experiments – carry multiple mutations.

Parallel evolution was common in our evolved phages, particularly in the control lines and the lineages that failed to infect their corresponding resistant *E. coli* C strains during the first evolution experiment (see **Table S3**). Of the three controls and 15 resistant lines that remained uninfected by their own phage, a total of 33 mutations were detected, but only ten were unique. All these mutations are located in the *F* gene, except one in the *A/A\** genes. *A* gene is involved in the stage II and III of phage DNA replication (Barrell et al. 1976; Sanger et al. 1978), while the *A\** gene is a non-essential gene that may play a role in the inhibition of host cell DNA replication and superinfection exclusion (Eisenberg and Ascarelli 1981). The mutation in the *F* gene at genomic position 2280 (causing amino acid change S427L in the F protein) arose in 13 lineages. Among them, nine (69 %) carry between one to three additional mutations. High degrees of parallel evolution have also previously been observed in ΦX174 wildtype strains and other viruses (Bull et al. 1997; Wichman et al. 1999; Remold et al. 2008; Bertels et al. 2019; Bertels et al. 2021) and is an indicator of either (i) a low number of mutational targets with similar high fitness gains (many more targets with lower fitness may exist but these will not be outcompeted by the high fitness mutants), or (ii) mutational bias towards certain positions in the genome (Lind et al. 2019; Bertels et al. 2021). While these substitutions may not allow the infection of resistant *E. coli* C strains, they are probably adaptations to our evolution experiment.

We also observed high levels of parallel evolution in the phage strains (isolates) that evolved to infect *E. coli* C mutants with the same predicted LPS phenotype (**Figs. 5A and 5B**). Parallelism was especially remarkable among phages infecting the four bacterial strains with the shortest (deep rough heptoseless) predicted LPS structure (see **Fig. 2**). The four phages infecting *E. coli* C R18, R8, R10, and R26 all carry an amino acid substitution at position 154 in the F protein (either Q154K or Q154R). In addition, the two phage strains adapted to the *E. coli* C mutants R8 (*hldE* mutant) and R18 (*waaC* mutant) also carry the mutations Y103H in the F protein and A68T in the H protein. Finally, while *E. coli* C R27 and R31 both became resistant via the same amino acid substitution in HldE (G27A), their corresponding phages evolved similarly by each acquiring a different mutation that led to the same change at the protein level (G91V in the H protein).

Parallel evolution was also observed in phages infecting bacteria within several other classes of predicted LPS structure. For example, the mutation at position 116 in the F protein (either D116G or D116S) occurred in four different phages all infecting different *waaO* mutants (**Figs. 5A and 5B**, **Table S2**). Parallel evolution in phages that evolved to infect *E. coli* C strains with the same predicted LPS structure generally supports the predicted classes of LPS structure. However, analyses in the following section demonstrate that LPS structure predictions generally underestimate LPS diversity.

### Evolved phages can be used to discriminate between LPS phenotypes

To determine precisely which of the 16 evolved phages can infect which of the 31 initially resistant *E. coli* C strains each (i.e., the infection pattern of each phage), a cross-infection assay was performed (**Fig. S3**). This involved measuring the infectivity of all possible host-phage combinations by two distinct methods: (i) phage spotting on host lawns and (ii) host spotting on phage lawns. In cases where a mismatch between the two results was observed, standard plaque assays were additionally applied (see **Methods**).

If our predictions – based on the current understanding in the literature – of seven distinct LPS structures among the 31 initially resistant *E. coli* C strains are accurate (see **Fig. 2 and Table S1**), then all phages that evolved to infect a mutant with a given predicted LPS structure are expected to cross-infect all other host strains with the same predicted LPS structure. That is, the infection patterns in **Fig. S3** would be identical for each class (colour) of *E. coli* C mutant.

To visually determine whether infection patterns cluster according to the predicted LPS structures, a hierarchical agglomerative clustering analysis was performed on the infection matrix in **Fig. S3**. Of the six predicted LPS structures that occur in at least two *E. coli* C mutants, three form unbroken clusters with respect to infectivity patterns (cyan, light purple, and dark purple highlighting in **Fig. 6**) and three do not (magenta, yellow, green highlighting). Each group is discussed in more detail below.

**Fig. 6.**
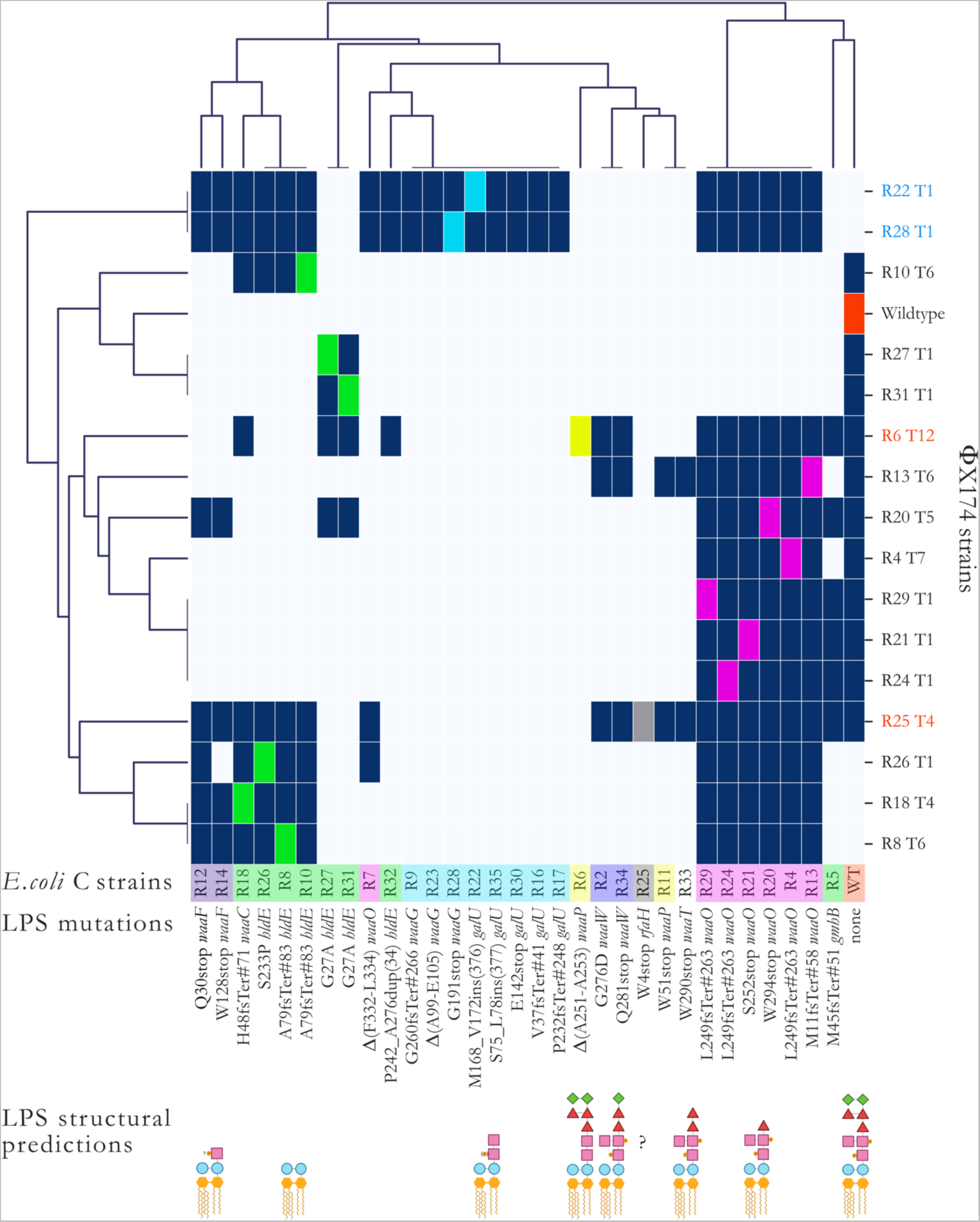
Hierarchical agglomerative clustering analysis of the host ranges of evolved phage. To unravel structures in the heatmap infection matrix (see **Fig. S3**), we performed a clustering analysis using the default settings of the Seaborn library (version 0.11.2) for Python (version 3.7.4). Briefly, each data point corresponds to either presence (dark blue) or absence (light blue) of phage infections determined from the combination of the different spotting test assays’ results (**Methods**). The linkage method computes the distance between them, then repeatedly combines the two nearest clusters into larger clusters until a single cluster is left. *E. coli* C strains are coloured based on their predicted core LPS structure (see **Fig. 2**). Phage names in black were obtained during the first 21 serial transfers (first evolution experiment). Phage strains in blue were obtained in the second evolution experiment (phage cocktail, one transfer per day). Phage strains in red were obtained in the third evolution experiment (phage cocktail, four transfers per day). Dark blue squares=infection, light blue square=no infection, coloured square=control infection by a phage evolved on that host. R# indicates the number of the resistant strain the phage evolved on, and T# is the transfer number where plaques were first observed. “?”: core LPS structure of *E. coli* C R25 (*rfaH* mutant) could not be predicted. “INS”: insertion. “DUP”: duplication. “stop”: stop codon. “Δ”: deletion. Examples: L249fsTer#263 indicates a frameshift (fs) leading to a premature codon stop (Ter); the position of the premature stop codon is in parentheses. S75_L78ins(377) indicates an insertion; the two flanking amino acids are separated by a “_” and followed by the number of inserted amino acids in parentheses. Δ(A99-E105) indicated deletion; two flanking amino acids are separated by a “-”. P242_A276dup(34) indicates a duplication; the two flanking amino acids are separated by a “_” and followed by the number of duplicated amino acids in parentheses.

*LPS predictions that cluster by infection pattern.* The identical ΦX174 mutants R22 T1 and R28 T1 that evolved to infect resistant strains with mutations in *galU* (*E. coli* C R22) and *waaG* (*E. coli* C R28) can cross-infect all eight *E. coli* C mutants predicted to exhibit the same LPS structure (*E. coli* C R9, R16, R17, R22, R23, R28, R30, R35; highlighted in cyan in **Figs. 6 and S3**). Furthermore, both *E. coli* C *waaW* mutants (R2 and R34; light purple) are cross-infected by the same three evolved phage genotypes (R6 T12, R13 T6, R25 T4). Also, both *E. coli* C *waaF* mutants (R12 and R14; dark purple) are infected by a near-identical set of seven (R12) or six (R14) phages. In each of these cases, phage infectivity patterns appear to be a good indicator of host LPS phenotype.

*LPS predictions that do not cluster by infection pattern.* Six of the seven *E. coli* C *waaO* mutants (highlighted in magenta in **Figs. 6 and S3**) can be cross-infected by the same set of 13 phages. The seventh *waaO* mutant (*E. coli* C R7) was the only *waaO* mutant for which the corresponding phage lineage did not evolve to infect the resistant mutant in the first evolution experiment. It can only be infected by the phage infecting R26 and phages that evolved in the second and third evolution experiments (**Fig. 6**). Hence, it is likely that the LPS structure of R7 differs from that of the other *waaO* mutants. The difference could conceivably result from residual (or altered) WaaO function; *E. coli* C R7 carries an in-frame deletion of a 6-bp repeat in *waaO*, which alters two WaaO residues but leaves the remainder of the protein intact (see **Table S1**). Because R7 is close to the *galU/waaG* mutants (**Figs. 6**), it may be possible that its LPS structure resembles the ones of the *galU/waaG* mutants. However, more detailed LPS studies are needed to elucidate the LPS structure of R7.

The infection patterns of the two *waaP* mutants *E. coli* C R6 and R11 (highlighted in yellow in **Figs. 6 and S3**) also suggest that, in contrast to the prediction, these strains possess different LPS structures. R6 is infected by a single phage that evolved directly on host R6 (phage R6 T12 from the third evolution experiment), while R11 is cross-infected by two different phages (phage R13 T6 from the first evolution experiment and phage R25 T4 from the third evolution experiment). *E. coli* C R6 was the last resistant strain to be infected by an evolved phage, and the evolved phage infecting R6 cannot cross-infect R11.

The R11 mutation introduces a premature codon stop at the beginning of the *waaP* gene (**Fig. 6 and Table S1**). *WaaP* is special because it is not required for completing the backbone structure of the LPS (Yethon et al. 1998). The deletion of this gene should, therefore, lead to a unique LPS structure (see **Fig. 2**). Instead, the cluster map analysis indicates that R11 displays an LPS phenotype comparable to the rough LPS phenotype of R33, a *waaT* mutant (**Fig. 6**). These similarities may be caused by a polar effect of the R11 mutation in *waaP* on *waaT* downstream. This effect may have resulted in a decrease or an alteration of *waaT* activity leading to a truncation of the outer core LPS similar to the one in R33.

The combination of phages that can infect each of the eight heptoseless *E. coli* C mutants predicted to have the shortest LPS structure (**Fig. 2**; green highlighting in **Figs. 6 and S3**) is remarkably different. These *E. coli* C strains cluster into four different phenotypic sub-groups (**Fig. 6**). The first sub-group consists of *E. coli* C R8, R10, R18, and R26. Each of these strains can be cross-infected by their corresponding evolved phages, regardless of their genotypes (a mixture of *waaC* or *hldE* mutants). However, none of the four evolved phages can cross-infect *E. coli* C R27 or R31 (*hldE* mutants). *E. coli* C R27 and R31 comprise the second sub-group. Both *E. coli* C R27 and R31 and their corresponding evolved phages acquired identical genotypes and, not surprisingly, they also share phenotypes. *E. coli* C R5 (*gmhB* mutant) and *E. coli* C R32 (*hldE* mutant) fall into a third and a fourth sub-groups. *E. coli* C R32 is the only heptoseless mutant for which the corresponding phage lineage failed to infect during the first evolution experiment. Only a very narrow but different set of evolved phages can infect R5 and R32 (**Figs. 6 and S3**).

A total loss of function in *hldE* or *waaC* genes always leads to a complete truncation of the inner core LPS (**Fig. S2A**). Thus, the involvement of these two genes in the production of the deep rough phenotypes (highlighted in green in **Figs. 6 and S3**) cannot explain the vast phenotypic diversity in LPS structures. For example, *E. coli* C R8, R10, R18, and R26 belong to the same phenotypic group despite having both *hldE* or *waaC* mutated, while R27, R31, and R32 (all *hldE* mutants) do not belong to this group (**Fig. 6 and Table S1**). Instead, the location and type of mutation are more likely to cause the observed LPS structure diversity.

Interestingly, HldE is a bifunctional protein, where each function is performed by a different domain: HldE1 (from M1 to T318) and HldE2 (from M344 to G477) (Valvano et al. 2000; Kneidinger et al. 2002; Raetz and Whitfield 2002; McArthur et al. 2005). Each domain has the potential to affect the LPS structure, but all four unique *hldE* mutations occur within the first domain only, resulting in at least three different LPS phenotypes (**Fig. 6 and Table S1**). These LPS phenotypes are caused by three different types of mutations (duplication, missense mutations, and frameshift) in different parts of the same domain. Yet, there is no obvious correlation between the LPS phenotypes and mutation positions and/or types. The phenotypic diversity produced by these mutations in a single domain of a single gene is surprising and highlights our extremely limited understanding of LPS biology.

The mutated *gmhB* gene in *E. coli* C R5 is involved in the production of yet another deep rough phenotype. Previous research demonstrated that the deletion of *gmhB* only causes a partial defect in the synthesis of the LPS core, resulting in the formation and co-existing of a heptoseless and a heptose-rich form of LPS molecules (Kneidinger et al. 2002). The *gmhB* mutation in R5 is an early frameshift leading to a premature stop codon. Hence, we assume that the effect is similar to what was found by Kneidinger et al., a mix of deep rough and wildtype LPS molecules on the outer bacterial cell membrane. This hypothesis is supported by the fact that R5 most closely clusters with the wildtype in our infection matrix.

### Linking phage genotype to phenotype

Phage infectivity patterns (host ranges) can be used to construct phage genotype-phenotype maps. For example, every phage strain that loses the ability to infect *E. coli* C wildtype has a combination of two mutations: A68T in the H protein and Q154K/R in the F protein (**Figs. 5B and 6**). In isolation, A68T confers the ability to infect *waaO* mutants and Q154K/R to infect [R8, R18, R10, R26]-heptoseless mutants. The two mutations combined allow the phage to infect both heptoseless and *waaO* mutants, but cannot infect the *E. coli* C wildtype (**Figs. 5B and 6**). ΦX174 R25 T4 is the only phage that can infect all three genotypes: *waaO*, the [R8, R18, R10, R26]-heptoseless mutants, and *E. coli* C wildtype. Presumably, because this phage does not contain the Q154K/R mutation and instead uses another set of mutations that allows R25 to infect [R8, R18, R10, R26]-heptoseless mutants (**Table S2**).

## Discussion

Our experiments have shown that *E. coli* C can become resistant to ΦX174 infection through mutations in genes involved in the LPS biosynthesis or assembly. ΦX174 can, in turn, evolve to reinfect initially resistant *E. coli* C strains by acquiring mutations in genes involved in host recognition or DNA injection. Similar to ΦX174 wildtype, the evolved phages are highly host-specific and hence can be used to distinguish between bacteria with different LPS structures. Such high host specificity allows testing of the currently accepted LPS structure model, where LPS structures are determined by the presence and absence of LPS synthesis and assembly genes. Our data on phage infectivity show that LPS diversity is far greater than the current model suggests. Presumably, the modulation of LPS gene activity leads to a variety of LPS phenotypes. Our findings can potentially be applied to other phages that use LPS as a single or complementary receptor for host infection (Bertozzi Silva et al. 2016; Latino et al. 2016; Broeker and Barbirz 2017). The LPS is a major structural component of the outer membrane of Gram-negative bacteria (Zhang et al. 2013), and its biosynthesis and assembly are conserved among bacterial species.

Another major finding of our study is that increasing phage and host diversity allows the infection of hard-resistant phenotypes (see **Fig. 4C**). This phenomenon – the successful infection of hard-resistant phenotypes – is similar to a receptor shift observed with phage λ (Meyer et al. 2012). Phage λ also requires at least four mutations to infect the novel OmpF receptor in *E. coli* B, making it almost impossible to evolve from the wildtype phage in a single step (Burmeister et al. 2016). Evolving all four mutations in one λ phage genome first required the presence of λ phage evolutionary intermediates that already carried specific mutations. These evolutionary intermediates may have arisen from the infection of easy-resistant *E. coli* B mutants (*malT* loss-of-function mutants) that could still spontaneously induce low levels of the traditional host receptor, LamB. By increasing the host diversity, phage λ eventually acquired additional mutations (four in total) to overcome the hard-resistant *E. coli B* mutants (*lamB* loss-of-function mutants) (Meyer et al. 2012; Gupta et al. 2022). Similarly, we showed that the successful infection of *waaG*, *galU*, *rfaH*, and *waaP*/*pssA* resistant strains became accessible by increasing host and phage diversity.

Our experiments suggest that it may be possible to breed well-established model phages such as ΦX174 to infect a wide range of currently resistant pathogenic *E. coli* strains. If possible, then this could make ΦX174 a promising therapeutic agent. Currently, instead of breeding well-known model systems to infect pathogenic bacteria, unknown phages that can infect specific bacterial pathogens are isolated from environmental samples (Chan et al. 2018). These unknown phages, however, still have to be characterized. Their safety has to be ensured by a process that, in the end, may drastically be more time-consuming than breeding well-studied model systems. Safety concerns regarding novel phages are significant due to their potential to carry dangerous toxins and antibiotic-resistance genes (Krüger and Lucchesi 2015; Colavecchio et al. 2017; Jamet et al. 2017). The considerable knowledge acquired from decades of research on ΦX174 structure, life cycle, and evolution showed that ΦX174 does not carry virulence genes. ΦX174 is also highly host-specific (Michel et al. 2010), meaning that it will be harmless to the patient’s microbiota, in contrast to phages with broader infectivity or antibiotics known to disrupt the microbiota and lead to adverse health outcomes (Ramirez et al. 2020). Furthermore, its use as a marker of immune responses in patients has already been approved *in vivo* by the FDA in specific case studies (Rubinstein et al. 2000; Bearden et al. 2005). ΦX174 can easily be manipulated in a laboratory unlocking its potential as a powerful therapeutic agent. For all these reasons, we believe that bacteriophage ΦX174 could become a promising therapeutic agent.

## Methods

### Bacterial and phage strains

The ancestral *Escherichia coli* strain C used in this study differs from the one uploaded on the NCBI website (GenBank accession number CP020543.1) by the presence of 9 additional insertions and two nucleotide substitutions (c → t at the position 1,1720,214 and g → t at the position 1,190,560). The genomic sequence of CP020543.1 was manually modified using Geneious Prime (version 2020.1.2). All resistant bacterial strains generated in this study are derived from our ancestral *E. coli* C strain. All bacteriophage strains are derived from the ancestral coliphage ΦX174 (GenBank accession number AF176034). AF176034 genomic sequence was downloaded from the NCBI website and manually annotated using Geneious Prime (version 2020.1.2). Whole-genome re-sequencing on ΦX174 wildtype’s glycerol stock showed no difference to the NCBI sequence. Complete genomes of *E. coli* C WT and ΦX174 WT used as references have been deposited in Zenodo (doi: 10.5281/zenodo.6952399). The *E. coli* C and ΦX174 ancestral strains were provided by Holly A. Wichman (Department of Biological Sciences, University of Idaho).

### Media and growth conditions

Phage and bacteria were grown in a shaking incubator (New Brunswick™ Innova^®^ 44) at 37°C, 250 rpm, in LB (Lysogeny Broth, Miller) medium supplemented with CaCl_2_ and MgCl_2_ at a final concentration of 5 mM and 10 mM, respectively. Solid LB agar (1.5 % agar) was used to plate both bacteria and phages. When plating phage, top agar overlay (0.5 % agar), also called <u>s</u>emi-<u>s</u>olid <u>a</u>gar (SSA) in this study, was supplemented with CaCl_2_ and MgCl_2_ at a final concentration of 5 mM and 10 mM respectively. PBJ solution (Phage Buffer Juice: 2.03 % Tris-HCl, 0.61 % MgCl_2_.6H_2_O) was used for serial dilutions. *E. coli* colony plates were generated by streaking with a sterile loop a scrap of the corresponding bacterial glycerol stock on the surface of a solid LB plate and incubated inverted overnight. After incubation, the colony plates were stored at 4°C for a maximum of two weeks. Overnight bacterial liquid cultures were systematically started by inoculating a single colony in 5 ml of LB supplemented with CaCl_2_ and MgCl_2_ at a final concentration of 5 and 10 mM, respectively, shaken at 250 rpm. Phage infections were either started from (i) a scrap of the corresponding phage glycerol stock resuspended in 100 μl of PBJ or (ii) a single plaque isolated from the top agar overlay with a sterile cut tip (a circle of agar) directly inoculated inside the bacterial culture. Overnight incubations of both plates and cultures were set up for ∼16-17 hours at 37°C.

### Measuring the growth of each resistant *E. coli* C strain

To assess the impact of LPS modifications harboured by the different *E. coli* C resistant mutants on their growth, two independent cultures of each resistant strain were grown at 37°C, 250 rpm for 17 hours. 180 μl of each overnight culture was inoculated into a 14 ml sterile tube (FALCON®) containing 5 ml of LB (supplemented with CaCl_2_ and MgCl_2_ at final concentrations of 5 and 10 mM, respectively) pre-warmed at 37°C. Incubation time was set to six hours at 37°C, 250 rpm. <u>O</u>ptical <u>d</u>ensity at 600 nm (OD600) was measured every 30 minutes with an Ultrospec 10 (Biochrom®). *E. coli* C wildtype was used as a reference. The mean OD600 values at each time point for each bacterial mutant were calculated and plotted.

### Isolation and storage of *E. coli* C strains with resistance to wildtype ΦX174

To avoid competition and to produce a diverse set of phage-resistant bacterial mutants with different growth phenotypes, we generated ΦX174-resistant *E. coli* C strains on agar plates. Thirty-five *E. coli* C wildtype colonies were randomly chosen from an agar plate, and each was used to inoculate an independent overnight liquid culture. 50 μl of each stationary phase culture was mixed with 50 μl of a high titer (∼10^9^ <u>p</u>laque-forming <u>u</u>nit pfu ml^-1^) stock of ΦX174 wildtype lysate, immediately plated with sterile beads on LB plates, and incubated at 37°C overnight. From each plate, half a randomly chosen colony was used to inoculate a fresh overnight LB liquid culture. Once grown, a sample from each culture was mixed with glycerol saline solution and frozen at -80°C (giving strains *E. coli* C R1-R35). To confirm the resistance to wildtype ΦX174, the remaining half of each colony was streaked onto an LB plate soaked with 100 μl of the high titer ΦX174 wildtype lysate and incubated at 37°C overnight.

### Phage lysate preparation

To extract phages in infected host cultures, ten to twelve drops of chloroform were added to growing *E. coli* C-phage co-cultures and vortexed for at least 30 seconds to kill the bacteria. Cultures were then centrifugated at 5000 rpm for 10 min at 4°C. Supernatants (phage lysates) were transferred to sterile 5 ml Eppendorf tubes and stored at 4°C. 1 ml of each lysate was stored with glycerol saline solution and frozen at -80°C.

### First phage evolution experiment - Standard

The protocol from Bono *et al*. (Bono et al. 2012) was adapted to evolve phages that can infect resistant *E. coli* C strains. Of R1-R35 – the 35 *E. coli* C strains resistant to ΦX174 wildtype, see above – 31 were used for the first phage evolution experiment (excluding R1, R3, R15, and R19; see **Results** and **Text S1**). Thirty-one ΦX174 lineages were each founded by ΦX174 wildtype and serially transferred daily, up to 21 days (21 transfers) in cultures containing a mix of *E. coli* C wildtype (permissive strain) and one resistant *E. coli* C strain (non-permissive strain; R2-R35). For each transfer, permissive and non-permissive bacteria were grown separately until they reached the log phase, then mixed in at a specific ratio depending on the growth of the non-permissive strain (see **Figs. 3A and S4** and **Supplementary Methods**) to a final volume of 5 ml. At each transfer, freshly prepared susceptible cells were infected by ΦX174 at MOI_input_ ∼0.1. Infected cultures were shaken at 250 rpm, 37°C, for three hours. Phage lysates were isolated as described in **Phage lysate preparation**. Phage lysates from the previous day were used to inoculate the fresh bacterial mixes the next day. Transfers continued until phages were found to infect their corresponding resistant *E. coli* C strain or until 21 transfers were reached. Three control lines were also run in parallel by transferring ΦX174 at MOI_input_ ∼0.1 in a culture containing *E. coli* C wildtype only. See **Supplementary Methods** for a detailed version of the protocol.

### Second phage evolution experiment – Increased diversity

A phage cocktail made of (i) the wildtype ΦX174 and (ii) all ΦX174 strains that successfully re-infected their corresponding resistant strains during the first phage evolution experiment was generated by first diluting each phage lysate and mixing them at a roughly equal number of <u>p</u>laque-forming <u>u</u>nits (PFU). Fourteen phage strains were used: ΦX174 R4 T7, ΦX174 R5 T2, ΦX174 R8 T6, ΦX174 R10 T6, ΦX174 R13 T6, ΦX174 R18 T4, ΦX174 R19 T1, ΦX174 R20 T5, ΦX174 R21 T1, ΦX174 R24 T1, ΦX174 R26 T1, ΦX174 R27 T1, ΦX174 R29 T1, and ΦX174 R31 T1. Phages infecting R5 and R19 were used for the cocktail but removed from the mutational analysis (see **Text S1 and Table S4**). The phage cocktail was grown and transferred daily for up to four days, on non-evolving host cultures containing (i) *E. coli* C wildtype, (ii) the 14 host strains (see **Text S1** and **Table S4**) for which resistance had been overcome (permissive hosts), and (iii) an excess of one of the still-resistant strain of interest (non-permissive host). For the non-permissive host, we used R6 (*waaP/pssA* mutant), R22 (*galU* mutant), R25 (*rfaH* mutant), or R28 (*waaG* mutant; see **Table S1**). *E. coli* C R22 and R28 were chosen as representatives of the *galU* and *waaG* mutants, respectively. Each bacterial strain (permissive and non-permissive) was grown separately in LB liquid culture for one hour (37°C, 250 rpm), then pooled at a roughly equal number of colony-forming units The resistant strain of interest was added in excess to give a final volume of 4 ml. Infected cultures were initiated by adding the phage cocktail (MOI_input_ ∼0.1), shaken at 250 rpm, 37°C, for 5 hours (see **Fig. 3B**). Phage lysates were isolated as described in **Phage lysate preparation**. Phage lysates from the previous day were used to inoculate the fresh bacterial mixes the next day. This process was repeated until phages were found to infect their corresponding resistant *E. coli* C strains or until four transfers had been completed. See **Supplementary Methods** for a detailed version of the protocol.

### Third evolution experiment – Increased diversity and generations

Protocol from the second evolution experiment was adjusted to reduce the amount of time necessary to retrieve the desired phages. The same phage cocktail used in the second phage evolution was grown and transferred daily, four times a day, for up to four days on non-evolving host cultures containing (i) *E. coli* C wildtype, (ii) the 14 host strains for which resistance had been overcome (permissive hosts), and (iii) one of the still-resistant strain of interest (non-permissive host). We increased the number of serial transfers per day from one to four, for a total of four days (16 transfers). *E. coli* C R6 (*waaP/pssA* mutant) and R25 (*rfaH* mutant, see **Table S1**) were used as non-permissive hosts. After reaching their exponential log phase, bacterial strains (except the non-permissive hosts) were mixed depending on their growth (**Figs. 3C and S4**, **Supplementary Methods**). New host cultures were kept in exponential growth phase independently by transferring every hour 1:5 of the volume (1 ml) in fresh LB pre-heated at 37°C (see **Fig. 3C**). Before each transfer, all host cultures were mixed together. Infected cultures were started by adding the phage cocktail to the final host mix at MOI_input_ ∼0.1, shaken at 250 rpm, 37°C, for one hour. After that, 1:15 of the volume of each infected culture was transferred in freshly mixed bacterial cultures and infection continued for one hour (see **Fig. 3C**). This step was repeated for a total of four transfers. The final (fourth) transfer lasted for two hours. All transfers completed on the same day involved transferring both phage and bacteria; on every fourth transfer, supernatants were collected and only phages were transferred. Phage lysates were isolated as described in **Phage lysate preparation**. Phage lysates from the previous day were used to inoculate the first fresh bacterial mixes the next day. This process was repeated until phages were found to infect their corresponding resistant *E. coli* C strains or until four transfers had been completed. See **Supplementary Methods** for a detailed version of the protocol.

### Isolation of evolved ΦX174 strains from evolution experiments

To isolate pure, single clones from the different evolution experiments, top agar overlays were prepared by mixing 100 μl of undiluted phage lysates with 200 μl of an overnight culture of the corresponding resistant *E. coli* C strain in 4 ml SSA. Plates were incubated inverted at 37°C for ∼16-17 hours. An isolated plaque was chosen randomly for each phage lysate to infect cultures of the corresponding resistant *E. coli* C strains (in exponential growth state) incubated at 37°C, 250 rpm, for 5 hours. Phage lysates were isolated as described in **Phage lysate preparation**. Phage isolates were re-isolated from single plaques a second time from their glycerol stocks. See **Supplementary Methods** for a detailed version of the protocol.

### Determination of the evolved phages’ host range by spotting assays

*Method 1.* Each top agar overlay was prepared by mixing 200 μl of an *E. coli* C strain from stationary phase culture in 4 ml SSA, then poured on LB plates and dried for at least 15 minutes. Then, 3 μl of each undiluted evolved phage lysate (between 10^7^ and 10^9^ pfu ml^-1^) was dropped at the surface. *Method 2*. Each top agar overlay was prepared by mixing a volume of each phage lysate in 4 ml SSA at a final concentration of ∼10^7^ pfu ml^-1^, then poured on LB plates and dried for at least 15 minutes. Then, 3 μl of both undiluted and ten-fold diluted of each *E. coli* C strain (from overnight culture) were dropped onto the surface with a pipette. For both methods, spots were dried for at least 30 minutes, and plates were incubated inverted at 37°C for ∼17 hours. *E. coli* C wildtype was used as a positive control (permissive strain), and *E. coli* K-12 MG1655 as a negative control (non-permissive strain). A bacterial host strain was classified as sensitive only when signs of lysis were detected using both methods in at least two (of three) replicates per method. A phage-bacterium combination that yielded different results between the two methods was tested in standard plaque assays. Finally, the host strain was classified as sensitive if plaques were observed at both dilutions. See **Supplementary Methods** for a detailed version of the protocol.

### Data visualisation

The hierarchical agglomerative clustering analysis (Clustermap **Fig. 6**), the heatmap (**Fig. S3**), and the growth curves (**Fig. S4**) were generated using the default settings of the Seaborn library (version 0.11.2) for Python (version 3.7.4). All schematic drawings (**Figs. 1**-**4**, **S1 and S2**) were created with <u>BioRender.com</u>. The Lollipop plot (**Fig. 5A**) was generated using trackViewer Vignette: lollipopPlot (Lolliplot) in R (version 1.27.11).

### Whole genome re-sequencing

*Bacteria.* Samples were prepared for whole genome re-sequencing from 1 ml of stationary-phase culture. Genomic DNA was extracted using the Wizard® Genomic DNA purification Kit (Promega, Germany). Extracted DNA was tested for quality, pooled, and sequenced by the Max-Planck Institute for Evolutionary Biology (Plön, Germany) using an Illumina Nextera DNA Flex Library Prep Kit to produce 150 bp paired-end reads (Picelli et al. 2014).

*Phage*. ΦX174 samples were prepared for whole genome re-sequencing from 1 ml of phage lysate. Genomic DNA was extracted using the QIAprep Spin Miniprep Kit (QIAGEN), then amplified by performing 20 cycles of PCR using Q5® High-Fidelity 2X Master Mix (NEB) (final concentration between 30-100 ng.µl^-1^). The primers used for the amplification of the ΦX174 genome are listed in **Table S6**. All PCR products were cleaned using the QIAquick PCR Purification Kit® (QIAGEN). DNA samples were tested for quality, pooled, and sequenced by the Max-Planck Institute for Evolutionary Biology (Plön, Germany). Sequencing was performed using an Illumina MiSeq DNA Flex Library Prep Kit to produce 150 bp paired-end reads (Picelli et al. 2014).

The quality of the sequencing output from bacteria and phages was controlled using *FastQC* version 0.11.8. Reads were trimmed using *Trimmomatic*, assembled and analysed using the *breseq* pipeline version 0.33.2 (Barrick et al. 2014; Deatherage and Barrick 2014; Deatherage et al. 2015) and Geneious Prime (version 2020.1.2).

### Sanger sequencing of ΦX174 *F* and *H* genes

Genomic DNA was extracted using the QIAprep Spin Miniprep Kit® (QIAGEN) (final concentration between 30-100 ng.µl^-1^). ΦX174’s *F* and *H* genes were amplified by performing 35 cycles of PCR using Phusion® High-Fidelity PCR Master Mix with HF Buffer (ThermoFisher). Primers used for this purpose are listed in **Table S6**. All PCR products were cleaned using the QIAquick PCR Purification Kit®. Sanger sequencing was performed using the sequencing primers listed in **Table S6**. The sequencing was performed at the Max-Planck Institute for Evolutionary Biology (Plön, Germany). Sequencing results were assembled and analysed with Geneious Prime (version 2020.1.2).

### Data availability

*Whole-genome and Sanger sequencing results.* Complete genomes of *E. coli* C WT and ΦX174 WT used as references and raw sequencing reads used to generate **Tables S1 to S5** have been deposited in Zenodo (doi: 10.5281/zenodo.6952399). *Growth curves.* Raw OD measurements used to generate **Fig. S4** have been deposited in Zenodo (doi: 10.5281/zenodo.6952399). *Hierarchical agglomerative clustering and heatmap data.* Pictures and raw data used to generate **Figs. 6** and **S3** have been deposited in Zenodo (doi: 10.5281/zenodo.6952399).

## Additional contact information

Jordan Romeyer Dherbey: – + 49 4522 763-278

Lavisha Parab: – + 49 4522 763-229

Jenna Gallie: – + 49 4522 763-574

Frederic Bertels: – + 49 4522 763-222

## Contributions

J.R.D and F.B conceived the study and planned the experiments. J.R.D and L.P. carried out the experiments. J.R.D conducted the sequence analyses. J.R.D, F.B, and J.G interpreted the results. J.R.D., F.B., and J.G. wrote the manuscript. J.R.D prepared all figures and tables. All authors reviewed and approved the manuscript.

## Acknowledgements

We received generous core funding from the Max Planck Society. We thank Loukas Theodosiou for his bioinformatic support, David Rogers for troubleshooting PCRs, and Sven Kuenzel for his help with phage and bacterial genome sequencing. We also thank the two reviewers for their time and insightful comments on the manuscript.

## Supplementary Text S1

*Excluded bacteria and phage isolates from downstream analyses*. We removed four bacterial strains (*E. coli* C R1, R3, R15, and R19) and two evolved phage strains (ΦX174 R5 T2 and ΦX174 R19 T1) from both mutational and phenotypical analyses. The reasons behind these exclusions are outlined below.

*Bacteria.* We found that both *E. coli* C R3 and R15’s glycerol stocks contain more than one genotype; while whole genome re-sequencing from their respective glycerol stocks showed only a single mutation in *galE*, whole genome re-sequencing of ten re-streaked colonies showed that additional mutations were systematically associated with the single mutation in *galE* (**Tables S4 and S5**). *E. coli* C R1 displayed an unstable resistant phenotype using the spotting assay *method 1* (see section **Determination of the evolved phages’ host range by spotting assays**). We performed phenotypic assays in a semi-solid environment by plating top agar overlays from overnight cultures of *E. coli* C R1 without phage. All overnight cultures were started with a randomly picked single colony obtained from the glycerol stock of R1. We observed that colonies of R1 made different lawn types: either “smooth” (no bacterial aggregate) or “granulous” (presence of numerous bacterial aggregates), which could potentially impact phage infectivity. The existence of two different aggregation phenotypes suggests that the glycerol stock of *E. coli* C R1 consists of two different populations. Unlike R3 and R15, however, no discrepancy was found between the whole genome re-sequencing results of R1 from the glycerol stock and its ten re-streaked colonies. The exact cause of the phenotypic inconsistency could not be identified. *E. coli* C R19 remains sensitive to ΦX174 wildtype infection. When plated undiluted, a high titer ΦX174 wildtype lysate (∼10^9^ pfu ml^-1^) could not clear *E. coli* C R19’s lawn, but a few thousand small clear plaques were produced. Thus, R19 is likely to be only partially resistant to ΦX174 wildtype infection. It carries a single mutation *yajC* (locus tag B6N50_17610), which encodes a periplasmic protein (Fang and Wei 2011) with a putative preprotein translocase subunit (Pfam e-value 2.3e-26) (see **Table S4**). While no link to the LPS biosynthesis or assembly has been defined yet, *yajC* could conceivably play a role in the injection of phage DNA into the bacterium’s cytoplasmic membrane (Schulze et al. 2014; Bohm et al. 2018).

In our matrices (**Figs. 6 and S3**), *E. coli* C R32 cannot be infected by any evolved phages from the first evolution experiment. The phage infecting R5 was the only phage that could infect R32 but was removed from the subsequent analysis (see below).

*Phage.* We removed the evolved phage infecting R5 (ΦX174 R5 T2) because its glycerol stock contains more than one genotype (confirmed by Sanger Sequencing). Since we removed R19 from the final analysis, we also removed its corresponding evolved phage obtained during the first evolution experiment (ΦX174 R19 T1). Details on their respective mutations (in the *H* gene) can be found in **Table S4**.

## Supplementary Figures

**Fig. S1.**
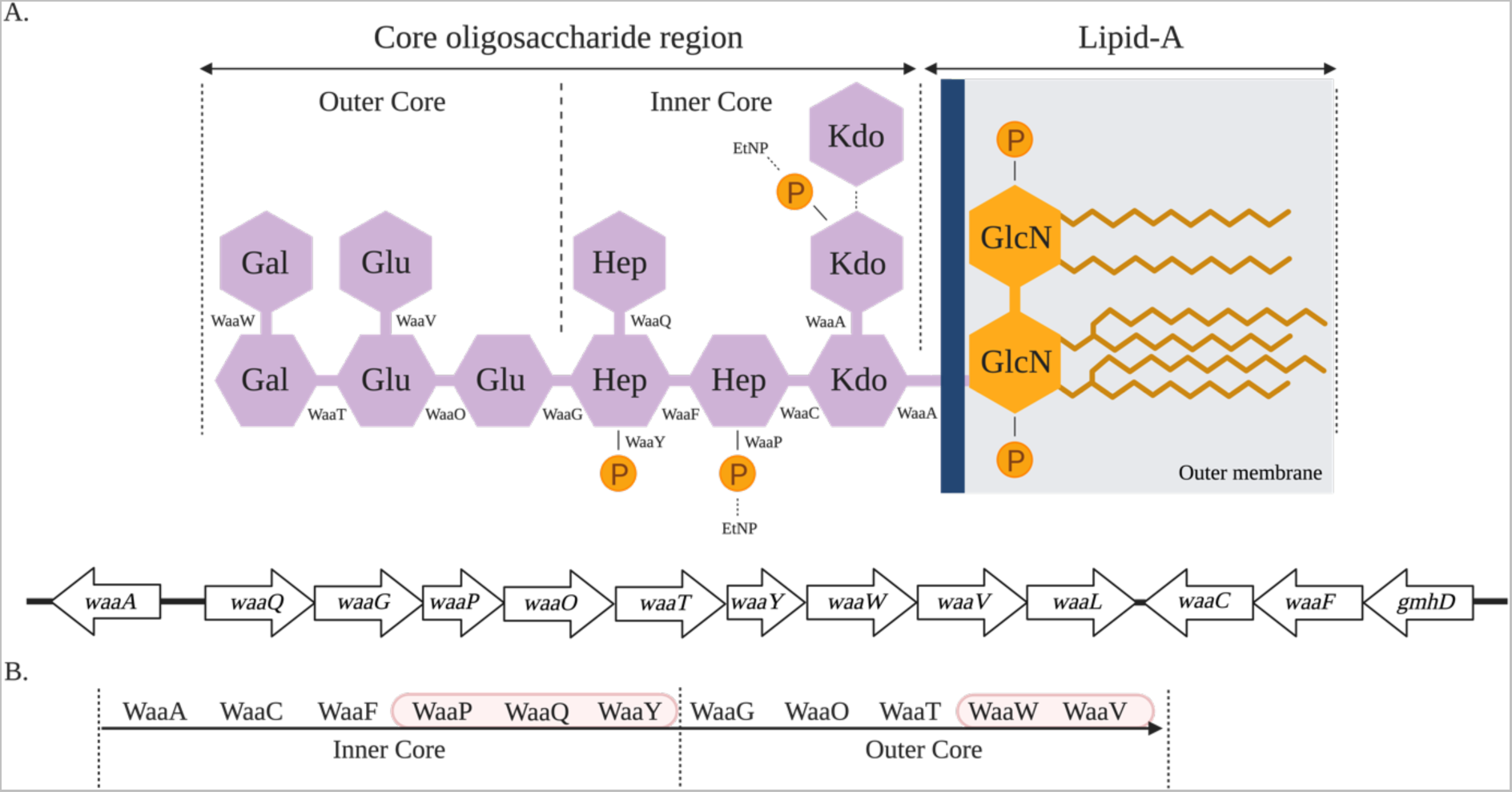
Assembly of *E. coli* rough type core LPS. **A.** Organization of the chromosomal *waa* region in the *E. coli C* wildtype strain. The chromosomal *waa* locus (formerly named *rfa*) is organized into three major operons, usually designated by the first gene of each transcriptional unit: *waaA, gmhD* and *waaQ* (Whitfield et al. 1999). The *waaA* operon is responsible for the incorporation of two Kdo (3-deoxy-D-*manno*-octulosonic acids) moieties to the lipid A (Belunis et al. 1995). The *gmhD* operon is required for the assembly of the inner core’s backbone (Schnaitman and Klena 1993), and the *waaQ* operon contains all eight genes necessary to modify the inner core and built the outer core (Whitfield et al. 1999) **B.** Sequential steps of the core OS LPS synthesis (from WaaA to WaaV). *Inner core assembly and modification*. Inner core assembly starts first with WaaA adding the two Kdo moieties to the lipid-A-IV precursor (Schnaitman and Klena 1993; Whitfield et al. 1999; Amor et al. 2000). A bacterium lacking WaaA activity displays severe membrane defects, resulting in the inability to form colonies (Klein et al. 2009). Heptose residues I and II are then anchored to the Kdo moieties by the heptosyltransferases WaaC and WaaF, respectively (Jansson et al. 1981; Schnaitman and Klena 1993; Whitfield et al. 1999). LPS core heptose kinases WaaP and WaaY add phosphate groups to the first and second heptose residues, respectively. WaaQ adds the last heptose to the second heptose residue (Jansson et al. 1981; Schnaitman and Klena 1993; Whitfield et al. 1999). *WaaP*, *waaQ*, and *waaY* proceed in this specific order. *WaaY* cannot work without the activity of *waaQ*, which cannot work without *waaP* (first red ellipse). *Outer core assembly*. The glucosyltransferases WaaG and WaaO start the formation of the outer core by anchoring the first glucose to the second heptose residue and the second glucose to the first glucose residue, respectively (Heinrichs, Yethon, and Whitfield 1998; Vinogradov et al. 1999; Whitfield et al. 1999). It has been suggested that the WaaG-catalyzed reaction might be required for WaaP and WaaY substrate specificity (Yethon et al. 2000). Then, the galactosyltransferases WaaT and WaaW add the first galactose to the second glucose residue and the second galactose to the first galactose residue, respectively (Heinrichs, Yethon, Amor, et al. 1998; Whitfield et al. 1999). The completion of the outer core LPS is achieved when WaaV adds the third glucose to the second glucose residue. Here, *waaV* requires the activity of *waaW* and proceeds in this specific order (second red ellipse). If *waaW* is deleted, *waaV* is not functional (Heinrichs, Yethon, Amor, et al. 1998; Leipold et al. 2007). Modifications of the LPS structure can lead to major phenotypic effects. In particular, the deep rough phenotype is usually associated with a strongly destabilized outer membrane, a decreased expression of some outer membrane proteins, a modification of the turgor pressure (Pagnout et al. 2019), and an increase of susceptibility to hydrophobic compounds such as AMPs (antimicrobial peptides), antibiotics, or bacteriocins (van der Ley et al. 1986; Schnaitman and Klena 1993; Yethon et al. 1998; Whitfield et al. 1999; Amor et al. 2000; Klein et al. 2013). Differences in LPS structures can also affect interactions with the host immune system (Raetz and Whitfield 2002; Matsuura 2013) and phage resistance (Hancock and Reeves 1976; Labrie et al. 2010; Kulikov et al. 2019; Mutalik et al. 2020).

**Fig S2.**
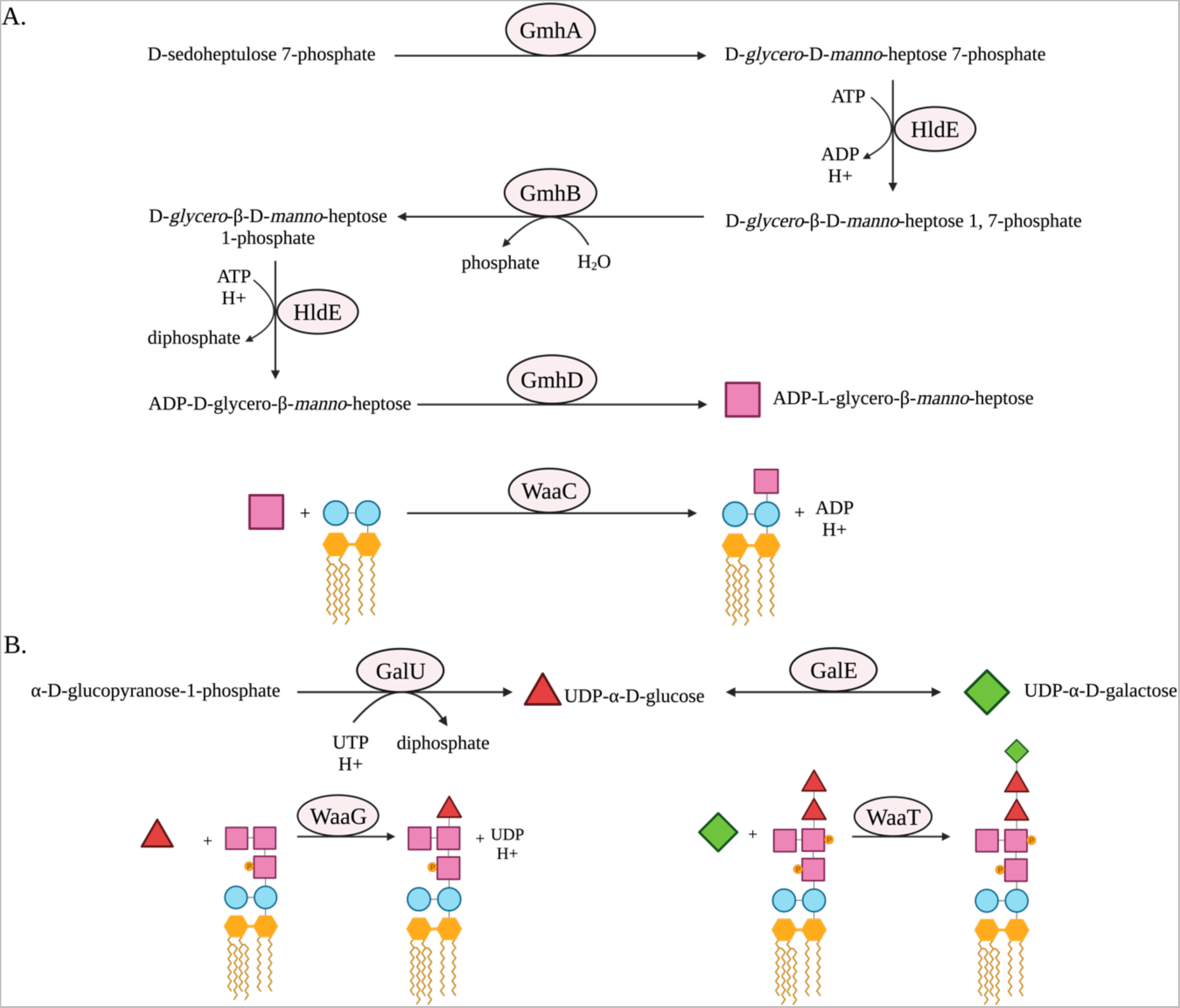
Biosynthesis of the core LPS glucose, galactose and heptose sugar components. **A.** ADP-L-glycero-β-D-*manno*-heptose synthesis pathway. It involves four genes: *gmhA*, *hldE*, *gmhB* and *gmhD*. Each gene encodes a protein that catalyses the production of a heptose intermediate. The final product is ADP-L-glycero-β-D-*manno*-heptose, which is used by the heptosyltransferase WaaC to build the inner core LPS (Kneidinger et al. 2002). Thus, deletion of one of these genes is expected to result in an heptoseless, deep rough LPS (McArthur et al. 2005). This figure is adapted from https://biocyc.org/ECOLI/NEW-IMAGE?type=PATHWAY&object=PWY0-1241 (Karp et al. 2019). **B.** Galactose degradation I (Leloir Pathway). GalU and GalE function in the Leloir pathway (Frey 1996; Kneidinger et al. 2002; McArthur et al. 2005). GalU catalyses the formation of UDP-α-D-glucose from α-D-glucopyranose 1-phosphate (Weissborn et al. 1994). UDP-α-D-glucose can then be incorporated into the outer core LPS by WaaG. Therefore, both Δ*galU* and Δ*waaG* mutants result in the same truncated LPS structure (Schnaitman and Klena 1993; Weissborn et al. 1994; Genevaux et al. 1999). GalE catalyses the interconversion of UDP-α-D-galactose and UDP-α-D-glucose during galactose catabolism (Pierson and Carlson 1996). UDP-α-D-galactose is then incorporated into the outer core LPS by WaaT (Heinrichs, Yethon, Amor, et al. 1998). Therefore, both Δ*galE* and Δ*waaT* are expected to result in the same truncated LPS (Schnaitman and Austin 1990). This figure was adapted from https://biocyc.org/ECOLI/NEW-IMAGE?type=PATHWAY&object=GALACTMETAB-PWY (Karp et al. 2019).

**Fig. S3.**
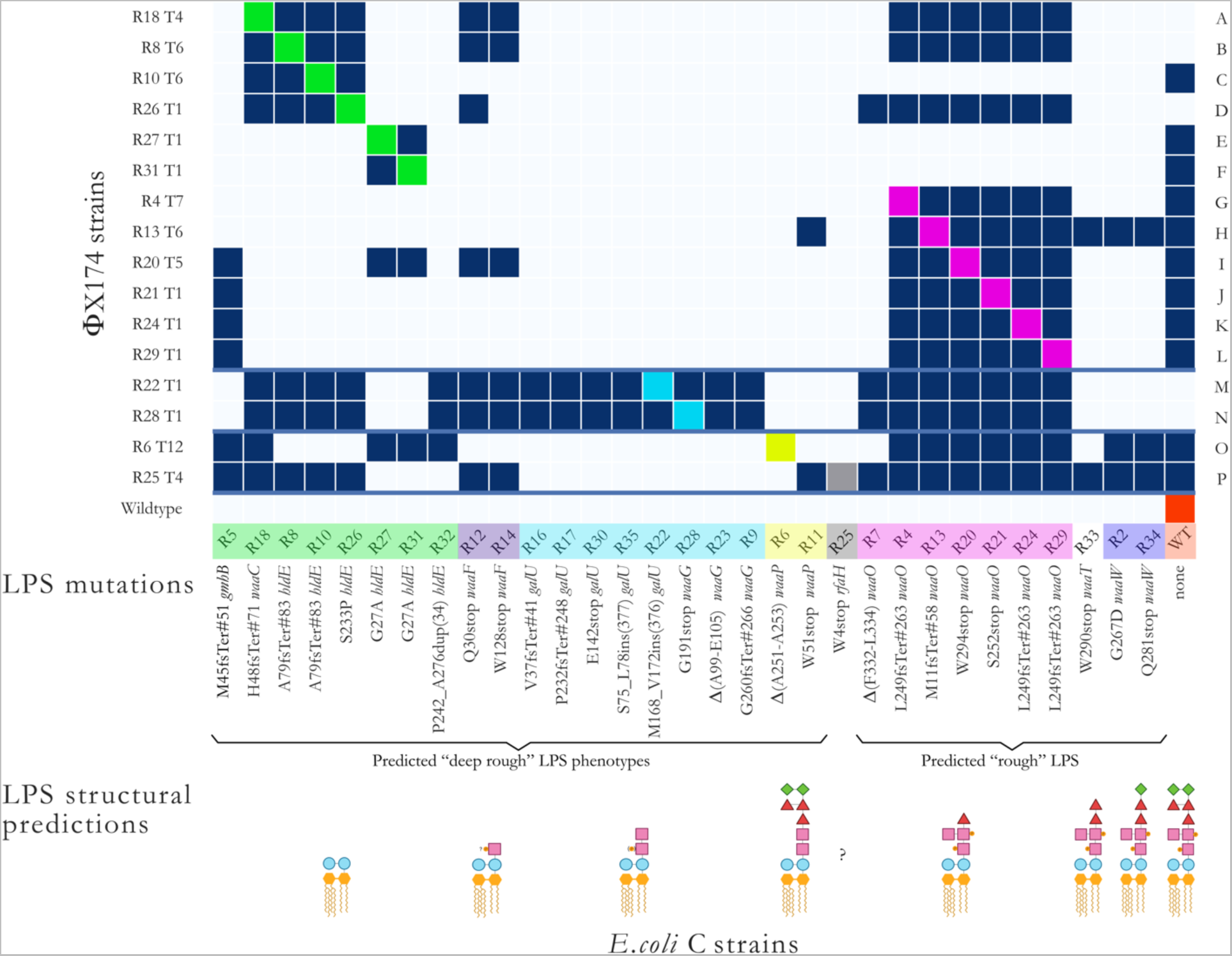
Infection matrix of evolved ΦX174 phages on the 31 resistant *E. coli* C strains. The infection matrix was produced by combining the results of two spotting methods and the plaque assays (see **Methods**). *E. coli* C R32 cannot be infected by any evolved phages from the first phage evolution experiment. The phage infecting R5 was the only phage that could infect R32 (**Text S1**) but was removed from the subsequent analysis due to a lack of isogeny*. E. coli* C strains are grouped and coloured based on their predicted core LPS structures (Jansson et al. 1981; Schnaitman and Austin 1990; Schnaitman and Klena 1993; Weissborn et al. 1994; Heinrichs, Yethon, and Whitfield 1998; Genevaux et al. 1999; Vinogradov et al. 1999; Whitfield et al. 1999; Amor et al. 2000; Kawaura et al. 2000; Yethon et al. 2000; Kneidinger et al. 2002; Raetz and Whitfield 2002; McArthur et al. 2005; Leipold et al. 2007; Fang and Wei 2011; Król et al. 2019). ΦX174 strains are ordered based on the predicted LPS structures they infected during the evolution experiments (A-F: overcame heptoseless *waaC* and *hldE* mutants; G-L: overcame *waaO* mutants; M-N: overcame *galU* and *waaG* mutants; O: overcame *waaP*/*pssA* mutant; P: overcame *rfaH* mutant). The solid blue lines separate the evolved phages according to the evolution experiment in which they were isolated (A-L: first; M-N: second; O-P: third). Dark blue squares=infection, light blue square=no infection, coloured square=control infection by a phage evolved on that host. R# indicates the number of the resistant strain the phage evolved on, and T# is the transfer number where plaques were first observed. “?”: core LPS structure of *E. coli* C R25 (*rfaH* mutant) could not be predicted. “INS”: insertion. “DUP”: duplication. “stop”: stop codon. “Δ”: deletion. Examples: L249fsTer#263 indicates a frameshift (fs) leading to a premature codon stop (Ter); the position of the premature stop codon is in parentheses. S75_L78ins(377) indicates an insertion; the two flanking amino acids are separated by a “_” and followed by the number of inserted amino acids in parentheses. Δ(A99-E105) indicated deletion; two flanking amino acids are separated by a “-”. P242_A276dup(34) indicates a duplication; the two flanking amino acids are separated by a “_” and followed by the number of duplicated amino acids in parentheses.

**Fig. S4.**
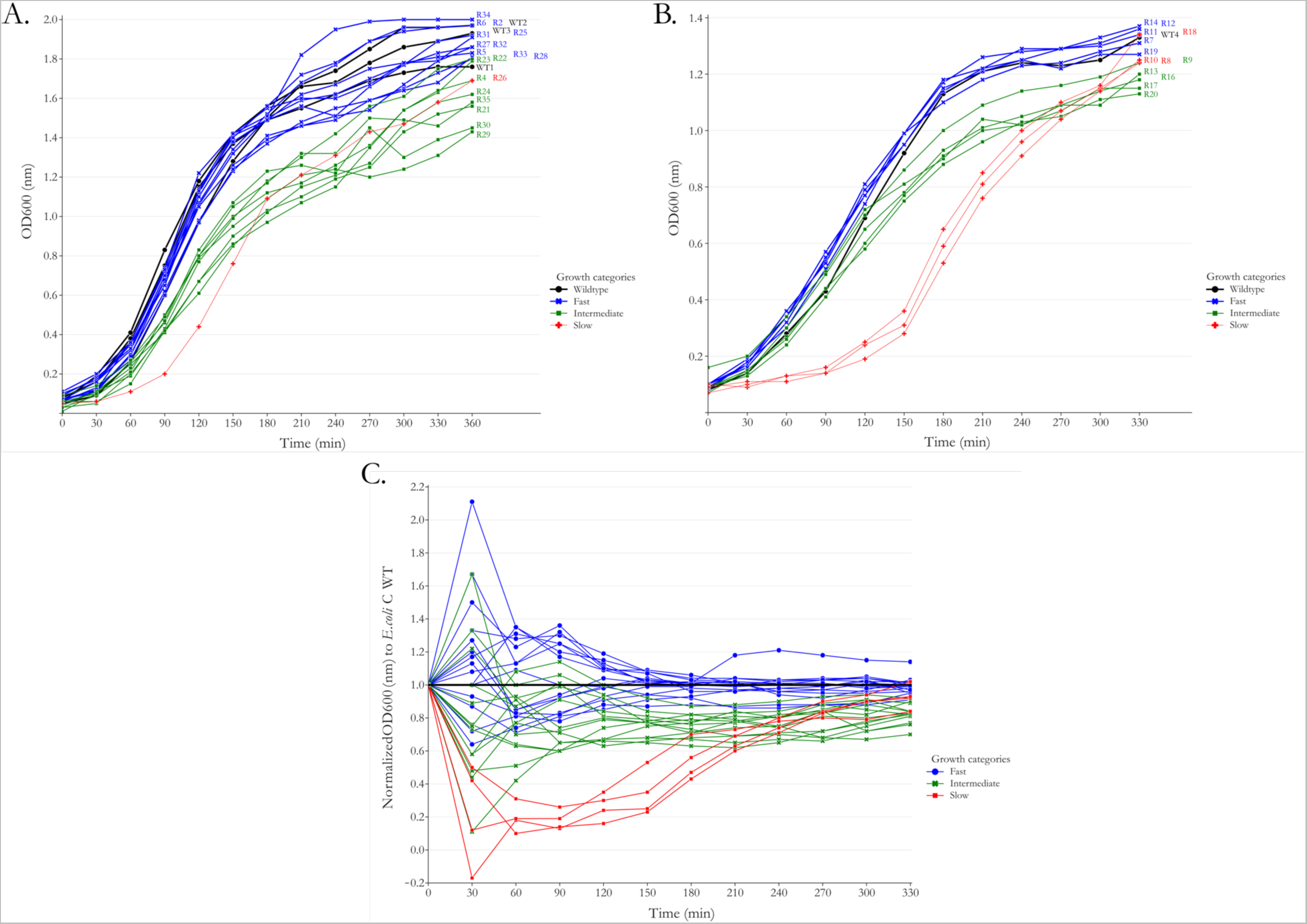
*E. coli* C resistant strains can be categorized as fast, intermediate, or slow growers. The mean OD600 values of two independent cultures grown in 5 ml LB at each time point for each bacterial mutant were calculated and plotted. Resistant strains are categorized with respect to their growth compared to *E. coli* C wildtype. Black lines: *E. coli* C wildtype. Blue lines: resistant bacteria that grew similarly to *E. coli* C wildtype (“fast growers”). Green lines: resistant strains grew somewhat more slowly than *E. coli* C wildtype and fast growers (“intermediate growers”). Red lines: resistant strains that grew more slowly than *E. coli* C wildtype and intermediate growers (“slow growers”). **A.** Growth curves of resistant strains R2-R6 and R21-R35. **B.** Growth curves of resistant strains R7-R20. These data (mutants and controls) were collected in a separate block to those in panel A. **C.** Mean OD600 values of all resistant strains in panels A and B, each normalized with respect to their corresponding *E. coli* C wildtype control ((OD600_R#_ti_ – ODR#_t0_)/(OD600_wildtype_ti_ – OD600_wildtype_t0_)). R# is the resistant strain number.

## Supplementary Tables

**Table S1:**
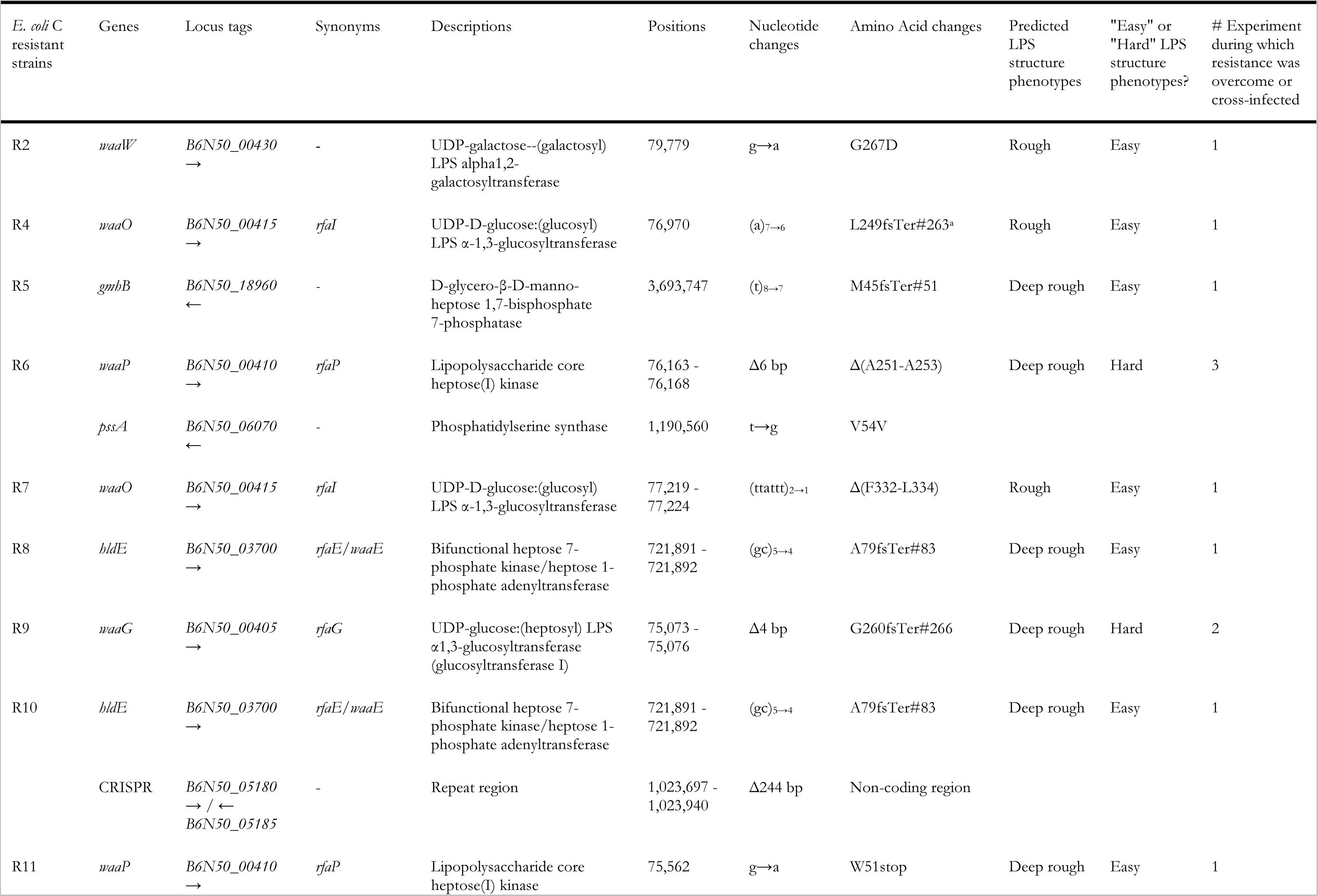
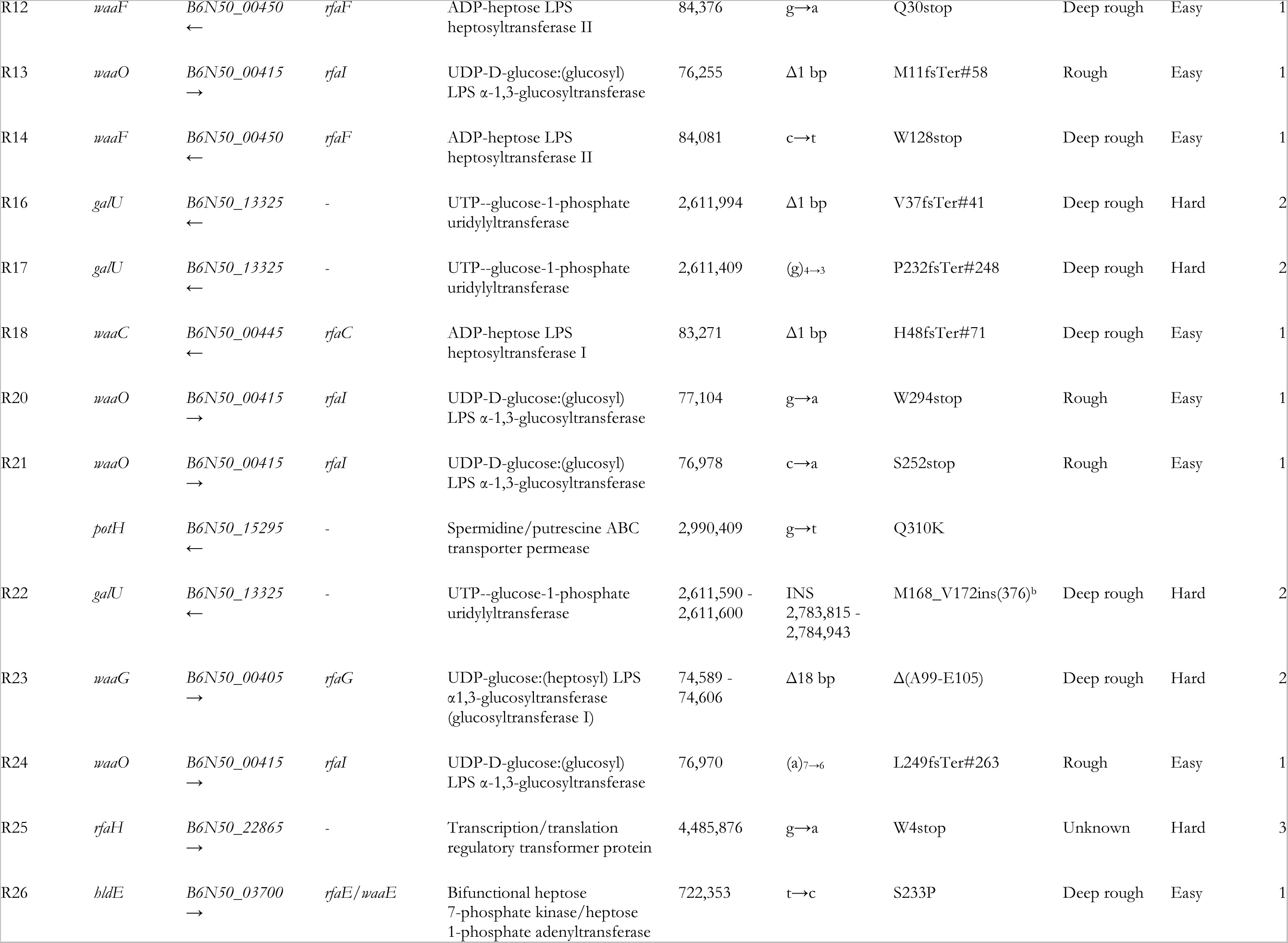
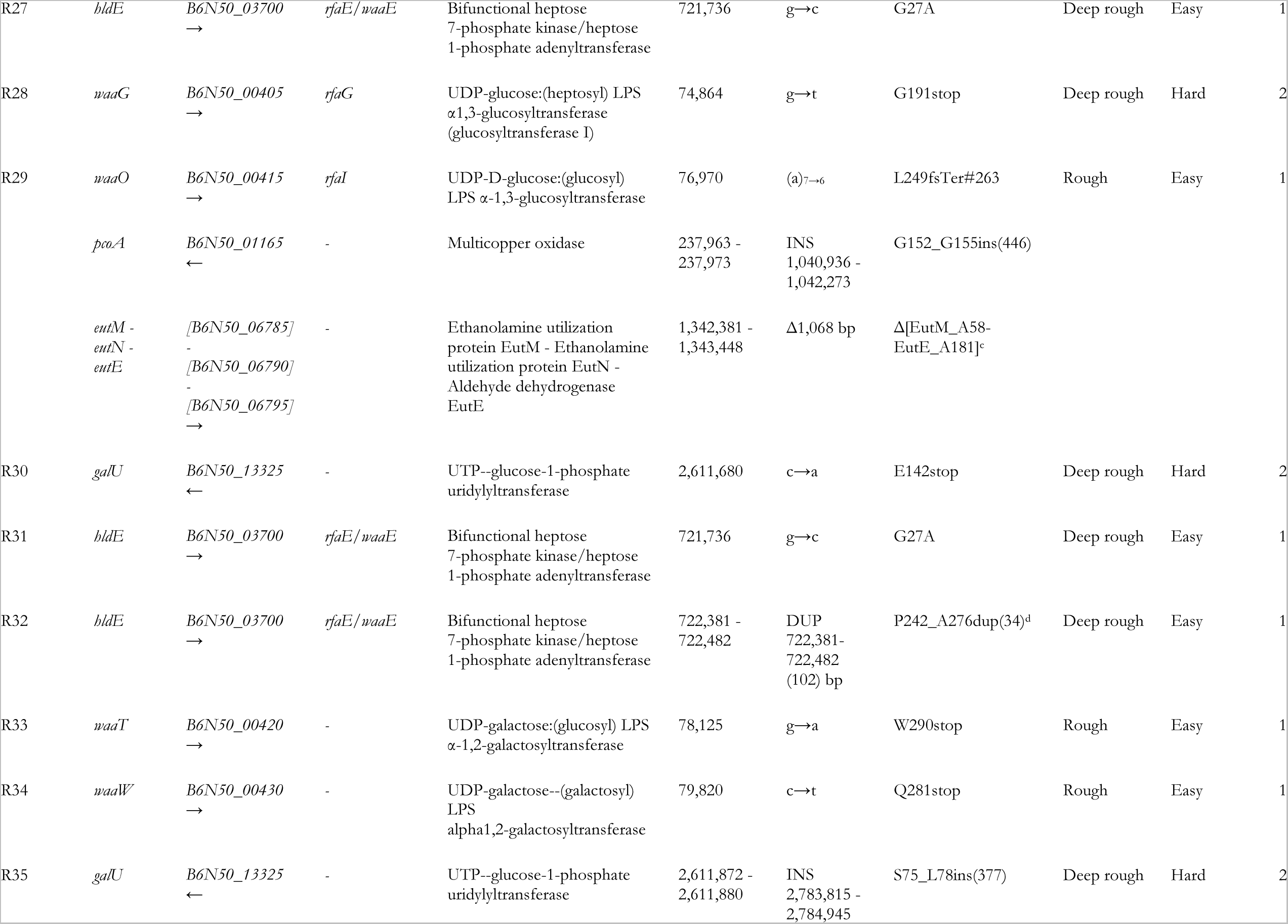
List of mutations found in *E. coli* C strains that are resistant to wildtype ΦX174. *E. coli* C resistant strains: R# is the strain number. Genes: name of the gene(s) in which mutations have been identified (as compared with wildtype *E. coli* C; doi: 10.5281/zenodo.6952399). Locus tags: identifier of each listed gene. Synonyms: alternative gene names. Descriptions: protein product encoded by each listed gene. Positions: genomic coordinates of mutation. Nucleotide changes: observed nucleotide change. Amino acid changes: resulting change in amino acid sequence. Predicted LPS structure phenotypes: based on predicted LPS structure (see **Fig. 2**). “Easy” or “Hard” LPS structure phenotypes?: easy LPS phenotypes were overcome or could be cross-infected after the first phage evolution experiment, while hard LPS phenotypes required the second or third phage evolution experiment to be infected. “INS”: insertion. “DUP”: duplication. “stop”: stop codon. “^a^”: L249fsTer#263 indicates a frameshift (fs) leading to a premature codon stop (Ter); the position of the premature stop codon is in parentheses. “^b^”: M168_V172ins(376) indicates an insertion; the two flanking amino acids are separated by a “_” and followed by the number of inserted amino acids in parentheses. “^c^”: Δ[EutM_A58-EutE_A181] indicated a long deletion starting from EutM to EutE. P242_A276dup(34) indicates a duplication; the two flanking amino acids are separated by a “_” and followed by the number of duplicated amino acids in parentheses.

**Table S2:** List of mutations identified in ΦX174 evolved isolates. ΦX174 strains: R# is the corresponding resistant strain number and T# is the transfer number where plaques were observed for the first time on the given resistant strain. Evolution experiment #: experiment number from which the evolved phages were isolated. Mutants sensitive to the evolved ΦX174 strains infection: list of all resistant strains that a specific phage strain can infect (See **Figs. 6 and S3**). Descriptions: names of proteins encoded by the listed genes. Positions: genomic coordinates of mutation (according to GenBank accession number AF176034.1). Nucleotide changes: observed nucleotide change. Amino acid changes: resulting change in amino acid sequence. Are mutations present in a control lineage?: indicates mutations that were observed in at least one control lineage (see **Methods**).

| ΦX174 strains | Evolution experiment # | Gene mutated in the corresponding *E. coli* C resistant strains | Mutants sensitive to the evolved ΦX174 strain infection | Gene mutated in ΦX174 strains | Descriptions | Positions | Nucleotide changes | Amino acid changes | Are mutations present in a control lineage? |
| --- | --- | --- | --- | --- | --- | --- | --- | --- | --- |
| R21 T1 | 1 | *waaO* | R5, R4, R13, R20, R21, R24, R29, WT | *F* | Capsid protein | 1,347 | a→g | D116G | No |
|  |  |  |  | *H* | Minor spike protein | 3,132 | g→a | A68T | No |
| R24 T1 | 1 | *waaO* | R5, R4, R13, R20, R21, R24, R29, WT | *F* | Capsid protein | 1,346 | g→a | D116S | No |
|  |  |  |  | *F* | Capsid protein | 1,347 | a→g | D116S | No |
| R26 T1 | 1 | *bldE* | R18, R8, R10, R26, R12, R7, R4, R13, R20, R21, R24, R29 | *F* | Capsid protein | 1,460 | c→a | Q154K | No |
|  |  |  |  | *H* | Minor spike protein | 3,132 | g→a | A68T | No |
| R27 T1 | 1 | *bldE* | R27, R31, WT | *H* | Minor spike protein | 3,202 | g→t | G91V | No |
| R29 T1 | 1 | *waaO* | R5, R4, R13, R20, R21, R24, R29, WT | *F* | Capsid protein | 1,346 | g→a | D116S | No |
|  |  |  |  | *F* | Capsid protein | 1,347 | a→g | D116S | No |
| R31 T1 | 1 | *bldE* | R27, R31, WT | *H* | Minor spike protein | 3,202 | g→t | G91V | No |
| R18 T4 | 1 | *waaC* | R18, R8, R10, R26, R12, R14, R4, R13, R20, R21, R24, R29 | *F* | Capsid protein | 1,307 | t→c | Y103H | No |
|  |  |  |  | *F* | Capsid protein | 1,461 | a→g | Q154R | No |
|  |  |  |  | *H* | Minor spike protein | 3,132 | g→a | A68T | No |
| R20 T5 | 1 | *waaO* | R5, R27, R31, R12, R14, R4, R13, R20, R21, R24, R29, WT | *F* | Capsid protein | 1,354 | t→a | N118K | No |
|  |  |  |  | *H* | Minor spike protein | 3,132 | g→a | A68T | No |
| R8 T6 | 1 | *bldE* | R18, R8, R10, R26, R12, R14, R4, R13, R20, R21, R24, R29 | *F* | Capsid protein | 1,307 | t→c | Y103H | No |
|  |  |  |  | *F* | Capsid protein | 1,461 | a→g | Q154R | No |
|  |  |  |  | *F* | Capsid protein | 3,132 | g→a | A68T | No |
| R10 T6 | 1 | *bldE* | R18, R8, R10, R26, WT | *F* | Capsid protein | 1,317 | a→g | H106R | No |
|  |  |  |  | *F* | Capsid protein | 1,460 | c→a | Q154K | No |
| R13 T6 | 1 | *waaO* | R11, R4, R13, R20, R21, R24, R29, R33, R2, R34, WT | *F* | Capsid protein | 2,015 | g→c | D339H | No |
| R4 T7 | 1 | *waaO* | R4, R13, R20, R21, R24, R29, WT | *F* | Capsid protein | 1,301 | a→g | T101A | Yes (in C1 T21) |
|  |  |  |  | *F* | Capsid protein | 1,347 | a→g | D116G | No |
|  |  |  |  | *F* | Capsid protein | 2,085 | c→t | A362V | No |
| R22 T1 | 2 | *galU* | R18, R8, R10, R26, R32, R12, R14, R16, R17, R30, R35, R22, R28, R23, R9, R7, R4, R13, R20, R21, R24, R29 | *F* | Capsid protein | 1,307 | t→c | Y103H | No |
|  |  |  |  | *F* | Capsid protein | 1,317 | a→g | H106R | No |
|  |  |  |  | *F* | Capsid protein | 1,461 | a→g | Q154R | No |
|  |  |  |  | *H* | Minor spike protein | 3,132 | g→a | A68T | No |
| R28 T1 | 2 | *waaG* | R18, R8, R10, R26, R32, R12, R14, R16, R17, R30, R35, R22, R28, R23, R9, R7, R4, R13, R20, R21, R24, R29 | *F* | Capsid protein | 1,307 | t→c | Y103H | No |
|  |  |  |  | *F* | Capsid protein | 1,317 | a→g | H106R | No |
|  |  |  |  | *F* | Capsid protein | 1,461 | a→g | Q154R | No |
|  |  |  |  | *H* | Minor spike protein | 3,132 | g→a | A68T | No |
| R25 T4 | 3 | *rfaH* | R5, R18, R8, R10, R26, R12, R14, R25, R11, R7, R4, R13, R20, R21, R24, R29, R33, R2, R34, WT | *F* | Capsid protein | 1,301 | a→g | T101A | Yes (in C1 T21) |
|  |  |  |  | *F* | Capsid protein | 1,304 | g→c | G102R | No |
|  |  |  |  | *F* | Capsid protein | 1,347 | a→g | D116G | No |
|  |  |  |  | *H* | Minor spike protein | 3,132 | g→a | A68T | No |
| R6 T12 | 3 | *waaP* | R5, R18, R27, R31, R32, R6, R4, R13, R20, R21, R24, R29, R2, R34, WT | *F* | Capsid protein | 1,727 | c→t | L243F | No |
|  |  |  |  | *F* | Capsid protein | 2,015 | g→c | D339H | No |
|  |  |  |  | *H* | Minor spike protein | 3,132 | g→a | A68T | No |
|  |  |  |  | *H* | Minor spike protein | 3,154 | c→t | A75V | No |

**Table S3:**
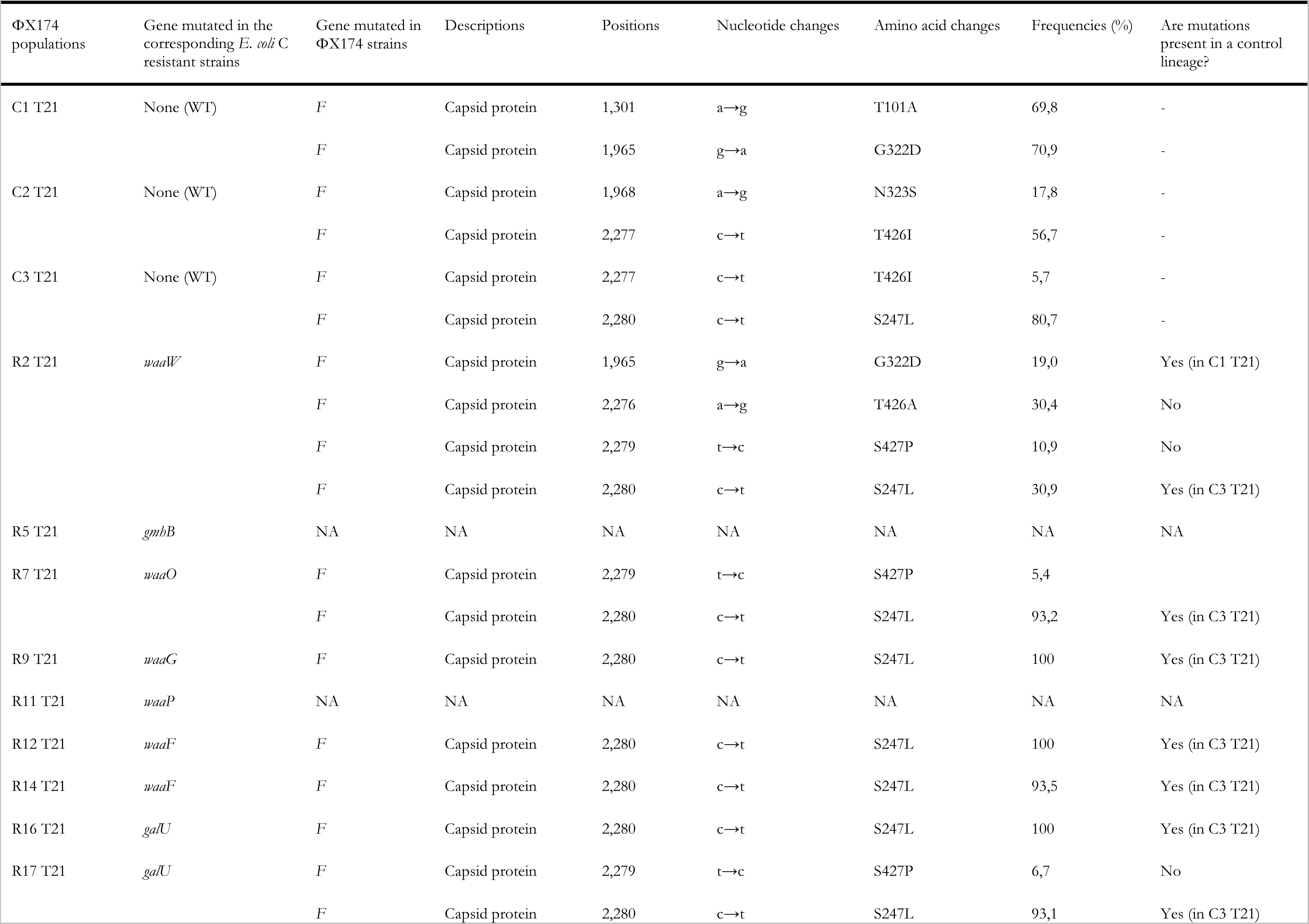
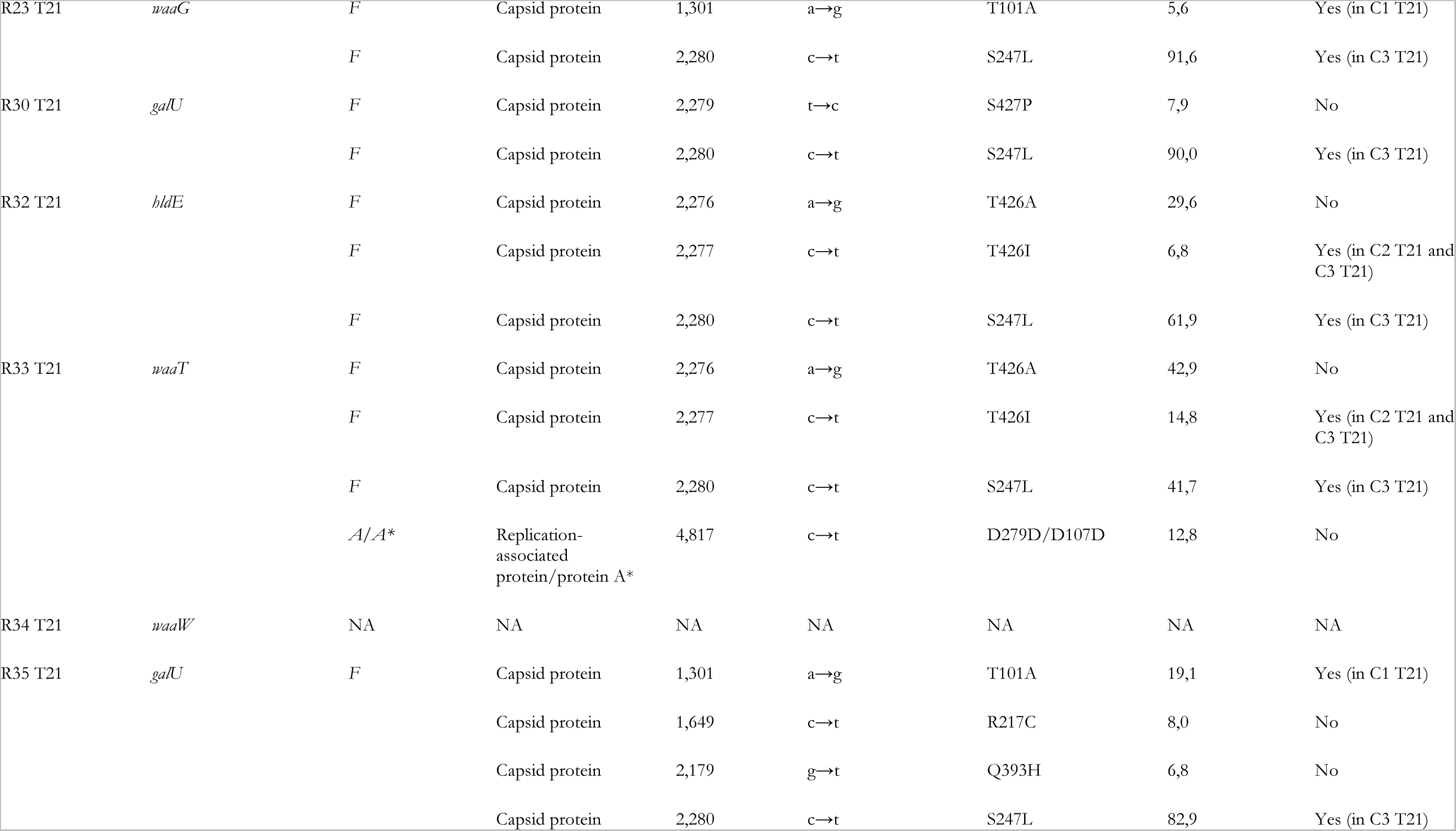
List of mutations found in the evolved ΦX174 populations that did not infect their corresponding *E. coli* C resistant strains. ΦX174 populations: R# is the corresponding resistant strain number and T# is the transfer number where plaques were observed for the first time on the given resistant strain. Descriptions: names of proteins encoded by the listed genes. Positions: genomic coordinates of mutation (according to GenBank accession number AF176034.1). Nucleotide changes: observed nucleotide change. Amino acid changes: resulting change in amino acid sequence. Frequencies (%): frequency of a given mutation in the phage population. Are mutations present in a control lineage?: indicates mutations that were observed in at least one control lineage (see Methods). NA: no data available.

**Table S4:**
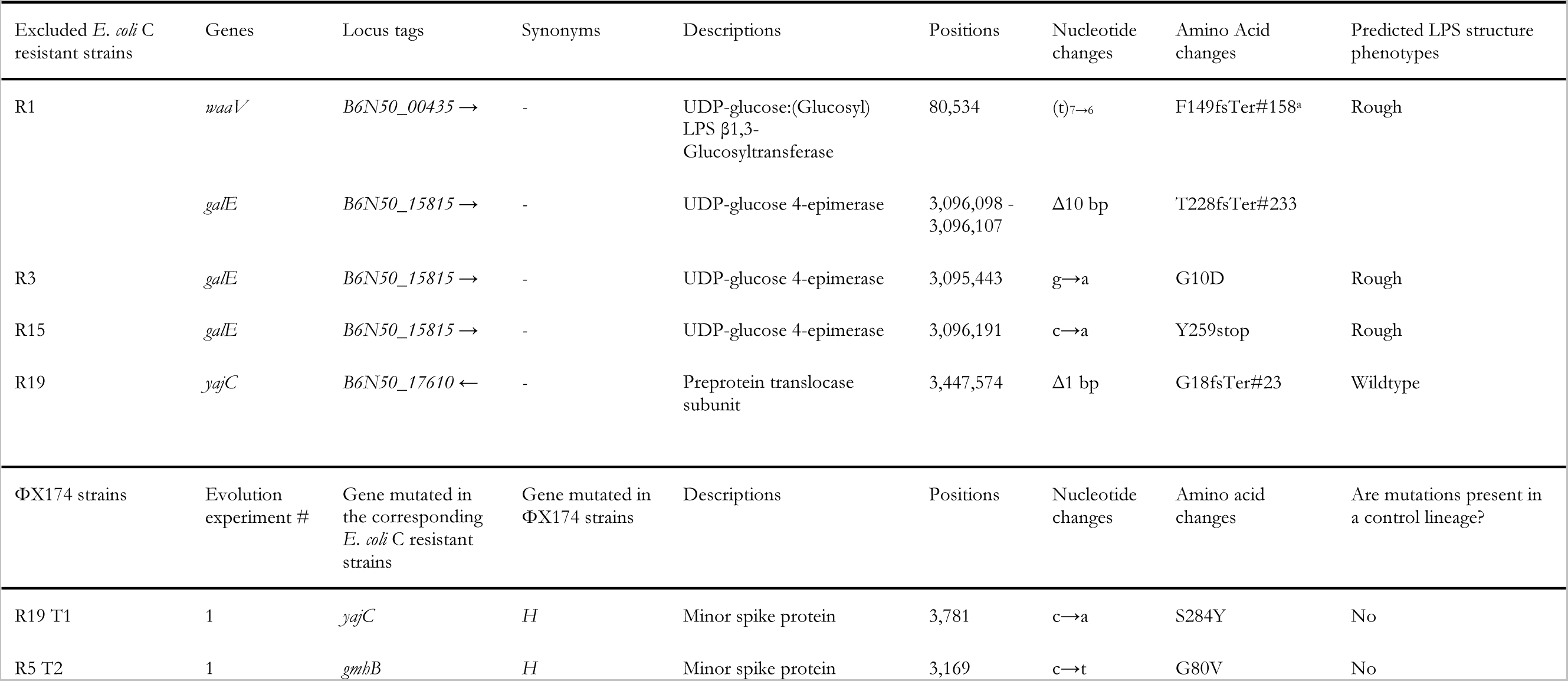
List of all *E. coli* C and evolved ΦX174 isolates excluded from the analysis. *E. coli* C resistant strains: R# is the excluded resistant strain number. Genes: name of gene(s) in which mutations have been identified (as compared with wildtype *E. coli* C; doi: 10.5281/zenodo.6952399). Locus tags: identifier of each listed gene. Synonyms: alternative gene names. Descriptions: protein product encoded by each listed gene. Positions: genomic coordinates of mutation. Nucleotide changes: observed nucleotide change. Amino acid changes: resulting change in amino acid sequence. Predicted LPS structure phenotypes: based on predicted LPS structure (see **Fig. 2**). “stop”: stop codon. “^a^”: F149fsTer#158 indicates a frameshift (fs) leading to a premature codon stop (Ter); the position of the premature stop codon is in parentheses. ΦX174 strains: R# is the corresponding resistant strain number and T# is the transfer number where plaques were observed for the first time on the given resistant strain. Evolution experiment #: experiment number where the evolved phages were obtained. Descriptions: names of proteins encoded by the listed genes. Positions: genomic coordinates of mutation (according to GenBank accession number AF176034.1). Nucleotide changes: observed nucleotide change. Amino acid changes: resulting change in amino acid sequence. Are mutations present in a control lineage?: indicates mutations that were observed in at least one control lineage (see **Methods**).

**Table S5:**
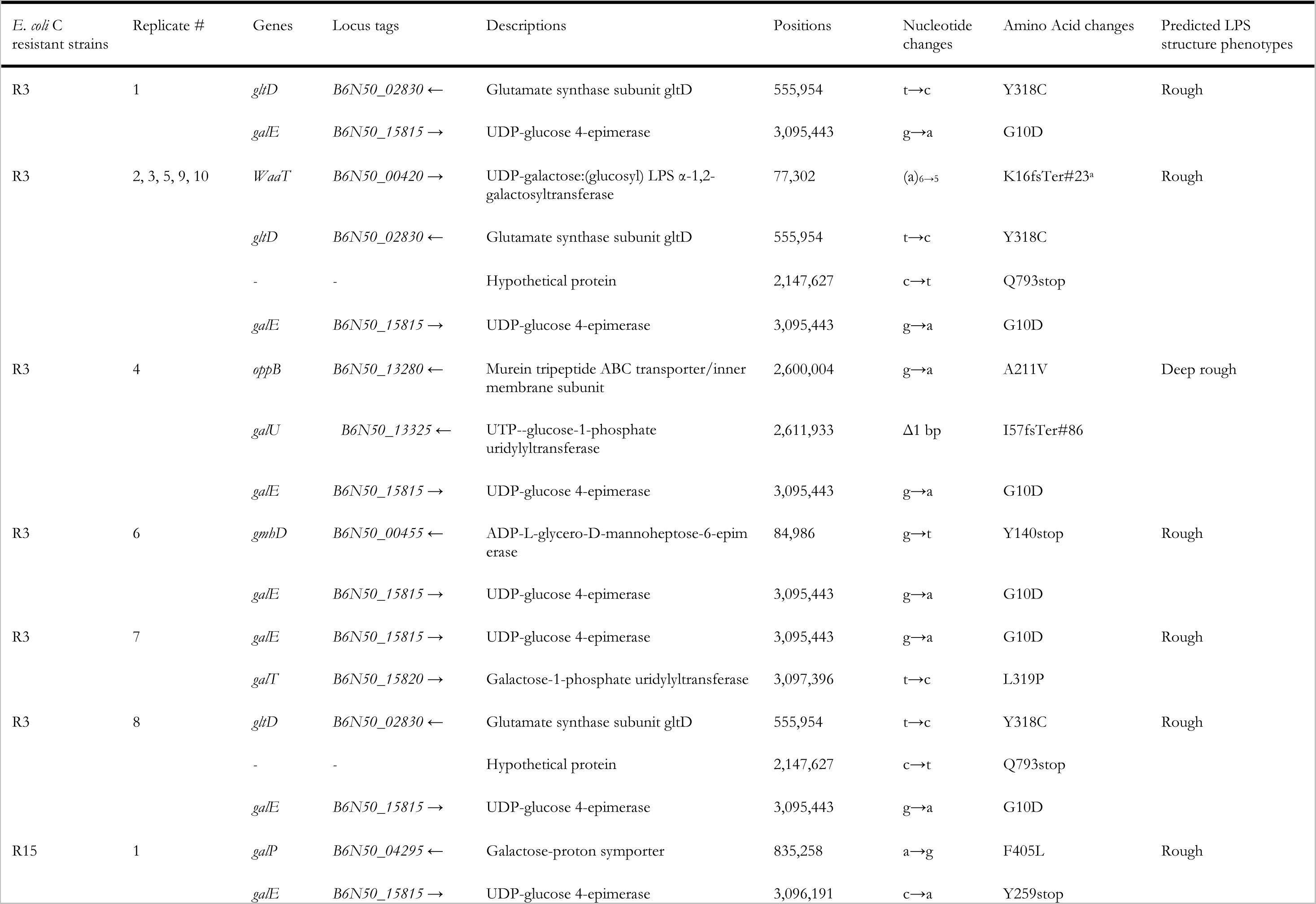
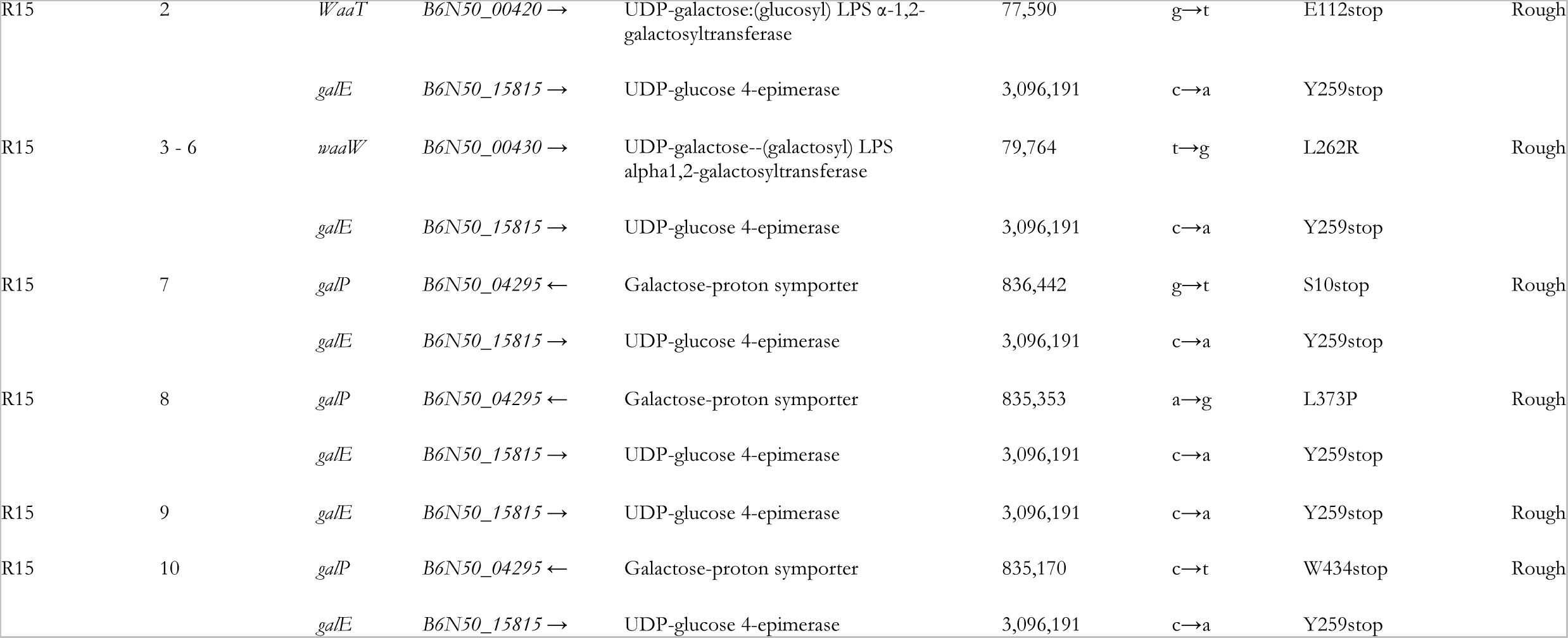
List of the additional mutations associated with *galE* mutations. *E. coli* C resistant strains: R# is the resistant strain number. Genes: name of the gene(s) in which mutations have been identified (as compared with wildtype *E. coli* C; doi: 10.5281/zenodo.6952399). Locus tags: identifier of each listed gene. Descriptions: protein product encoded by each listed gene. Positions: genomic coordinates of mutation. Nucleotide changes: observed nucleotide change. Amino acid changes: resulting change in amino acid sequence. Predicted LPS structure phenotypes: based on predicted LPS structure (see **Fig. 2**). “stop”: stop codon. “^a^”: K16sTer#23 indicates a frameshift (fs) leading to a premature codon stop (Ter); the position of the premature stop codon is in parentheses.

**Table S6.**
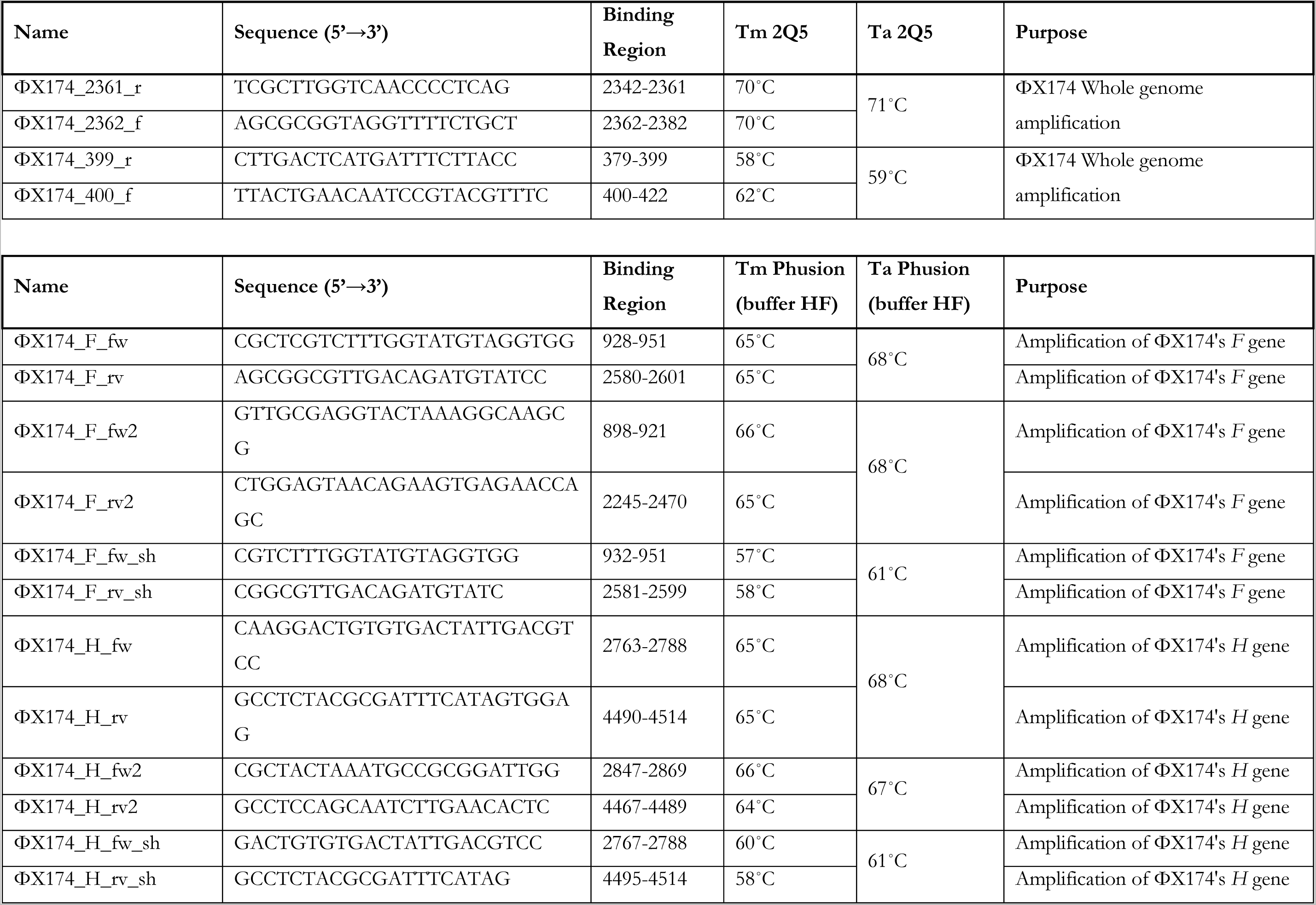
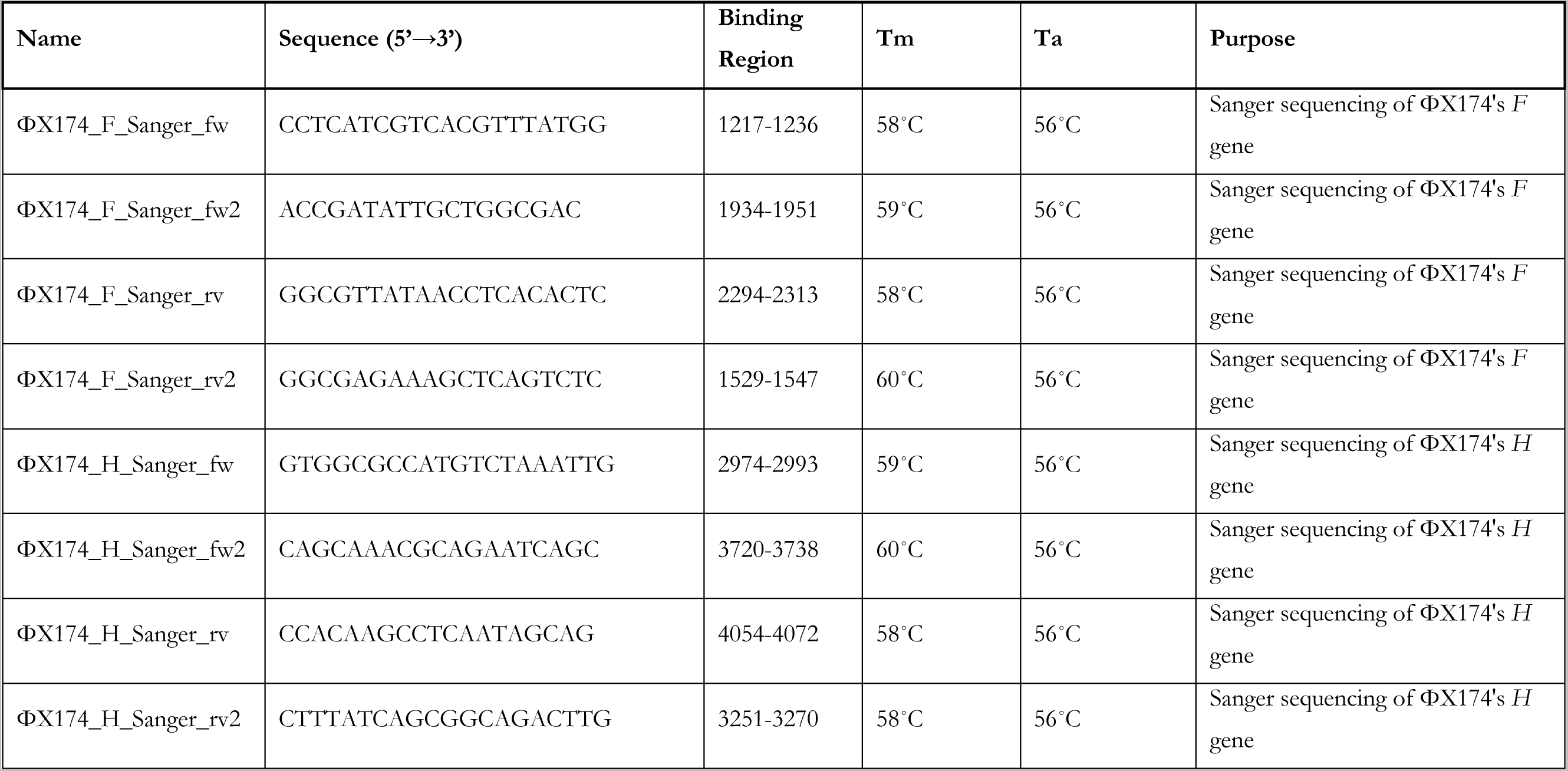
List of primers used in this study. *F*: encodes for the viral capsid protein. *H*: encodes for the minor spike protein. “fw”: forward. “rv”: reverse. Tm 2Q5: melting temperature of the primer used with the Q5® High-Fidelity 2X Master Mix (NEB). Ta 2Q5: annealing temperature of the set of primers used with the Q5® High-Fidelity 2X Master Mix (NEB). Tm Phusion (buffer HF): melting temperature of the primer used with Phusion® High-Fidelity PCR Master Mix with HF Buffer (ThermoFisher). Ta Phusion (buffer HF): annealing temperature of the primer used with Phusion® High-Fidelity PCR Master Mix with HF Buffer (ThermoFisher).

## Supplementary Methods

In this section, we provide a more detailed version of the protocols used in the **Methods** section.

### First phage evolution experiment - Standard

The protocol from Bono *et al*. (Bono et al. 2012) was adapted to evolve phages that can infect resistant *E. coli* C strains. Of R1-R35 – the 35 *E. coli* C strains resistant to ΦX174 wildtype, see above – 31 were used for the first phage evolution experiment (excluding R1, R3, R15, and R19; see **Results** and **Text S1**). Thirty-one ΦX174 lineages were each founded by ΦX174 wildtype and serially transferred daily, for up to 21 days, on non-evolving host cultures containing a mix of (i) *E. coli* C wildtype (permissive strain) and (ii) one resistant *E. coli* C strain (non-permissive strain; R2-R35).

For each transfer, permissive and non-permissive bacteria were grown separately until they reached the log phase. These cultures were founded with an initial inoculum of 180 μl from fast and intermediate growers’ overnight cultures or 350 μl from slow growers’ overnight cultures (see **Fig. S4**). Fast and intermediate growers’ cultures were incubated at 37°C (250 rpm) for two hours, while slow growers’ cultures were incubated for three hours. Then, permissive and non-permissive strains were mixed in a 14 ml sterile tube at a specific ratio depending on the growth of the non-permissive strain (1:1 for fast growers, 1:2 for intermediate growers, 1:3 for slow growers), to a final volume of 5 ml.

At each transfer, ∼10^8^ freshly prepared susceptible cells were infected by ∼10^7^ ΦX174 (MOI_input_ ∼0.1). Infected cultures were shaken at 250 rpm, 37°C for three hours. Phage lysates were isolated as described in **Phage lysate preparation**. Phage lysates from the previous day were used to inoculate the fresh bacterial mixes the next day. Transfers continued until phages were found to infect their corresponding resistant *E. coli* C strain, or until 21 transfers were reached. Three control lines were also run in parallel, by transferring ΦX174 at MOI_input_ ∼0.1 in a culture containing *E. coli* C wildtype (permissive strain) only.

### Second phage evolution experiment – Increased diversity

A phage cocktail made of (i) the wildtype ΦX174 and (ii) all ΦX174 strains that successfully re-infected their corresponding resistant strains during the phage evolution experiment – Standard was generated by first diluting each phage lysate and mixing them at a roughly equal number of pfu (final concentration of ∼2.10^8^ pfu ml^-1^). These phage strains (14 in total) are: ΦX174 R4 T7, ΦX174 R5 T2, ΦX174 R8 T6, ΦX174 R10 T6, ΦX174 R13 T6, ΦX174 R18 T4, ΦX174 R19 T1, ΦX174 R20 T5, ΦX174 R21 T1, ΦX174 R24 T1, ΦX174 R26 T1, ΦX174 R27 T1, ΦX174 R29 T1, and ΦX174 R31 T1. Despite being removed from the mutational analysis, phages infecting R5 and R19, respectively, were part of the final cocktail (see **Text S1** and **Table S4**).

The phage cocktail was grown and transferred daily, for up to four days, on non-evolving host cultures containing: (i) *E. coli* C wildtype, (ii) the 14 host strains for which resistance had been overcome (permissive hosts: *E. coli* C R4, R5, R8, R10, R13, R18, R19, R20, R21, R24, R26, R27, R29, and R31), and (iii) an excess of one of the still-resistant strain of interest (non-permissive host). For the non-permissive host, we used R6 (*waaP/pssA* mutant), R22 (*galU* mutant), R25 (*rfaH* mutant), or R28 (*waaG* mutant; see **Table S1**). *E. coli* C R22 and R28 were chosen as representatives of the *galU* and *waaG* mutants, respectively.

Each bacterial strain (permissive and non-permissive) was grown separately in LB liquid culture for one hour (37°C, 250 rpm). An inoculum of 180 μl from overnight cultures was used for the fast and intermediate growers, while 350 μl was used for the slow growers. After the incubation time, permissive strains were pooled at a roughly equal number of <u>c</u>olony-forming <u>u</u>nits (CFU), for a final concentration of ∼10^8^ susceptible, permissive cells. One of the *E. coli* C resistant strains of interest (non-permissive host; R6, R22, R25 or R28) was then added in excess (in between ∼1 and 5*x*10^8^ resistant cells) in the host mix, to give a final volume of 4 ml.

Infections were initiated by adding ∼10^7^ phages (MOI_input_ ∼0.1) from the phage cocktail to the bacterial culture mix. Infected cultures were shaken in 100 ml sterile Erlenmeyer at 250 rpm, 37°C, for five hours. Phage lysates were isolated as described in **Phage lysate preparation**. Phage lysates from the previous day were used to inoculate the fresh bacterial mixes the next day. This process was repeated until phages were found to infect their corresponding resistant *E. coli* C strains, or until four transfers had been completed.

### Third evolution experiment – Increased diversity and generations

The same phage cocktail used in the second phage evolution experiment – Increased diversity was grown and transferred daily, four times a day, for up to four days on non-evolving host cultures containing (i) *E. coli* C wildtype, (ii) the 14 host strains for which resistance had been overcome (permissive hosts: *E. coli* C R4, R5, R8, R10, R13, R18, R19, R20, R21, R24, R26, R27, R29, and R31), and (iii) one of the still-resistant strain of interest (non-permissive host). The definitive protocol from the second phage evolution experiment – Increased diversity was adjusted to reduce the amount of time necessary to retrieve the desired phages. We increased the number of transfers per day from one to four for a total of four days (16 transfers in total). Only *E. coli* C R6 (*waaP/pssA* mutant) and R25 (*rfaH* mutant) (see **Table S1**) were used as non-permissive hosts.

Each bacterial strain (permissive and non-permissive) was grown separately for one hour. An inoculum of 180 μl from overnight cultures was used for the fast and intermediate growers, while 350 μl was used for the slow growers. After the incubation time, permissive strains were pooled at a roughly equal number of <u>c</u>olony-forming <u>u</u>nits (CFU) in three different mixes depending on the growth categories of each bacterium (slow, intermediate, and fast growers, **Fig. S4**). 1 ml of each fast grower was used to make the “fast mix” (constituted of *E. coli* C wildtype, R5, R19, R27, and R31). 833 μl of each intermediate grower was used for the “intermediate mix” (constituted of R4, R13, R20, R21, R24, and R29). Finally, 1.25 ml of each slow grower was used to prepare the “slow mix” (constituted of R8, R10, R18, and R26).

Non-permissive strains and bacterial mixes were kept in exponential growth phase independently by transferring 1:5 of the volume (1 ml) in fresh LB pre-heated at 37°C every hour (final volume of 5 ml). To check whether the growth of the different bacteria remained consistent from one transfer to another, the OD600 values were measured before each transfer. The volume transferred at each hour was modified accordingly if the bacterial growth was too fast or too slow.

Before adding the phage cocktail, all bacterial mixes were pooled together at a roughly equal number of cfu (500 μl each, for a total of ∼10^8^ susceptible cells), and one of the *E. coli* C resistant strains of interest (non-permissive host; R6 or R25) was added in excess (i.e., the number of resistant cells was comprised between ∼1 and 5.10^8^), for a final volume of 4 ml.

Infections started by adding ∼10^7^ phages (MOI_input_ ∼0.1) from the cocktail in the final bacterial mix. Infected cultures were shaken in 100 ml sterile Erlenmeyer at 250 rpm, 37°C, for one hour. After that, 1:15 of the volume of each infected culture was transferred to freshly mixed bacterial cultures and the infection continued for one hour. This step was repeated for a total of four transfers. The final (fourth) transfer lasted for two hours to ensure that all phages completed the infection cycle and are present in the culture medium and not in the cell. All transfers completed on the same day involved transferring both phage and bacteria; on every fourth transfer, supernatants were collected and only phages were transferred. Phage lysates were isolated as described in **Phage lysate preparation**. Phage lysates from the previous day were used to inoculate the first fresh bacterial mixes the next day. This entire process was repeated until phages that could infect their corresponding resistant *E. coli* C strains were identified or until four transfers had been completed.

### Isolation of evolved ΦX174 strains from single plaques

To isolate pure, single clones from the different evolution experiments, top agar overlays were prepared by mixing 100 μl of undiluted phage lysates with 200 μl of an overnight culture of the corresponding resistant *E. coli* C strain in 4 ml SSA (supplemented with CaCl_2_ and MgCl_2_ at a final concentration of 5 and 10 mM, respectively). Top agar overlays were poured onto LB plates, dried for at least 15 minutes, and incubated inverted at 37°C for ∼16-17 hours. Plaques were counted the next day. An isolated plaque was chosen randomly for each phage lysate to infect cultures of the corresponding resistant *E. coli* C strains (in exponential growth state) incubated at 37°C, 250 rpm, for five hours. Phage lysates were isolated as described in **Phage lysate preparation**. To obtain pure phage glycerol stocks (isogenic stocks which are presumed to hold one genetic clone (van Charante et al. 2019)), phages were purified via a second round of plaque isolation from their first glycerol stocks. Phage lysates were isolated and stored as described in **Phage lysate preparation**.

### Determination of the evolved phages’ host range by spotting assays

*Method 1.* Each top agar overlay was prepared by mixing 200 μl of an *E. coli* C strain from stationary phase culture in 4 ml SSA (supplemented with CaCl_2_ and MgCl_2_ at a final concentration of 5 and 10 mM, respectively), then poured on LB plates and dried for at least 15 minutes under a sterile laminar flow hood. Then, 3 μl of each undiluted evolved phage lysate (between 10^7^ and 10^9^ pfu ml^-1^) was dropped at the surface.

*Method 2*. Each top agar overlay was prepared by mixing a volume of each phage lysate in 4 ml SSA at a final concentration of ∼10^7^ pfu ml^-1^, then poured on LB plates and dried for at least 15 minutes under a sterile laminar flow hood. Then, 3 μl of both undiluted and ten-fold diluted of each *E. coli* C strain (from overnight culture) were dropped onto the surface with a pipette.

For both methods, spots were dried for at least 30 minutes under sterile laminar flow, and the plates were subsequently incubated inverted at 37°C for ∼17 hours. *E. coli* C wildtype was used as a positive control (permissive strain), and *E. coli* K-12 MG1655 as a negative control (non-permissive strain). Sterile H_2_O was also spotted at the end of each plate as a control for material contamination (pipettes and tips). A bacterial host strain was classified as sensitive only when signs of lysis were detected using both methods, in at least two (of three) replicates per method. A phage-bacterium combination that yielded different results between the two methods was tested in *standard plaque assays* as follows: a top agar overlay was prepared by mixing 100 μl of the phage lysate diluted to 10^-1^ and 10^-6^ with 200 μl of the resistant *E. coli* C strain from stationary phase culture in 4 ml SSA supplemented with CaCl_2_ and MgCl_2_ at a final concentration of 5 and 10 mM, respectively), then poured on LB plates and dried for at least 15 minutes. The presence or absence of plaques was assessed the next day. Finally, the host strain was classified as sensitive if plaques were observed at both dilutions.

## References

1. Amor K, Heinrichs DE, Frirdich E, Ziebell K, Johnson RP, Whitfield C. 2000. Distribution of core oligosaccharide types in lipopolysaccharides from *Escherichia coli*. Infect Immun. 68:1116–1124.

2. Bailey MJA, Hughes C, Koronakis V. 1997. RfaH and the *ops* element, components of a novel system controlling bacterial transcription elongation. Mol Microbiol. 26:845–851.

3. Barrell BG, Air GM, Hutchison CA. 1976. Overlapping genes in bacteriophage ΦX174. Nature 264:34–41.

4. Barrett RDH, Schluter D. 2008. Adaptation from standing genetic variation. Trends Ecol Evol. 23:38– 44.

5. Barrick JE, Colburn G, Deatherage DE, Traverse CC, Strand MD, Borges JJ, Knoester DB, Reba A, Meyer AG. 2014. Identifying structural variation in haploid microbial genomes from short-read resequencing data using *breseq*. BMC Genomics. 15:1039.

6. Bearden CM, Agarwal A, Book BK, Vieira CA, Sidner RA, Ochs HD, Young M, Pescovitz MD. 2005. Rituximab inhibits the *in vivo* primary and secondary antibody response to a neoantigen, bacteriophage ΦX174. Am J Transplant. 5:50–57.

7. Bertels F, Leemann C, Metzner KJ, Regoes RR. 2019. Parallel evolution of HIV-1 in a long-term experiment. Mol Biol Evol. 36:2400–2414.

8. Bertels F, Metzner KJ, Regoes R. 2021. Convergent evolution as an indicator for selection during acute HIV-1 infection. Peer Community J. 1:e4.

9. Bertozzi Silva J, Storms Z, Sauvageau D. 2016. Host receptors for bacteriophage adsorption. FEMS Microbiol Lett. 363:fnw002.

10. Bono LM, Gensel CL, Pfennig DW, Burch CL. 2012. Competition and the origins of novelty: experimental evolution of niche-width expansion in a virus. Biol Lett. 9:20120616–20120616.

11. Broeker NK, Barbirz S. 2017. Not a barrier but a key: how bacteriophages exploit host’s O-antigen as an essential receptor to initiate infection. Mol Microbiol. 105:353–357.

12. Bull JJ, Badgett MR, Wichman HA, Huelsenbeck JP, Hillis DM, Gulati A, Molineux IJ. 1997. Exceptional convergent evolution in a virus. Genetics. 147:1497–1507.

13. Bull JJ, Levin BR, Molineux IJ. 2019. Promises and pitfalls of *in vivo* evolution to improve phage therapy. Viruses. 11:1083.

14. Burmeister AR, Lenski RE, Meyer JR. 2016. Host coevolution alters the adaptive landscape of a virus. Proc R Soc B Biol Sci. 283:20161528.

15. Centers for Disease Control and Prevention (U.S.). 2019. Antibiotic resistance threats in the United States, 2019. Available from: https://stacks.cdc.gov/view/cdc/82532.

16. Chan BK, Turner PE, Kim S, Mojibian HR, Elefteriades JA, Narayan D. 2018. Phage treatment of an aortic graft infected with *Pseudomonas aeruginosa*. Evol Med Public Health. 2018:60–66.

17. Colavecchio A, Cadieux B, Lo A, Goodridge LD. 2017. Bacteriophages contribute to the spread of antibiotic resistance genes among foodborne pathogens of the *Enterobacteriaceae* family – A review. Front Microbiol. 8:1108.

18. Cox J, Putonti C. 2010. Mechanisms responsible for a ΦX174 mutant’s ability to infect *Escherichia coli* by phosphorylation. J Virol. 84:4860–4863.

19. Crill WD, Wichman HA, Bull JJ. 2000. Evolutionary reversals during viral adaptation to alternating hosts. Genetics. 154:27–37.

20. Deatherage DE, Barrick JE. 2014. Identification of mutations in laboratory-evolved microbes from next-generation sequencing data using *breseq*. Methods Mol Biol. 1151:165–188.

21. Deatherage DE, Traverse CC, Wolf LN, Barrick JE. 2015. Detecting rare structural variation in evolving microbial populations from new sequence junctions using *breseq*. Front Genet. 5:468.

22. Eisenberg S, Ascarelli R. 1981. The A* protein of ΦX174 is an inhibitor of DNA replication. Nucleic Acids Res. 9:1991–2002.

23. Feige, Ulrich, Stirm, Stephan. 1976. On the structure of *Escherichia coli* C cell wall lipopolysaccharide core and its ΦX174 receptor region. Biochem Biophys Res Commun. 71:8.

24. Gupta A, Zaman L, Strobel HM, Gallie J, Burmeister AR, Kerr B, Tamar ES, Kishony R, Meyer JR. 2022. Host-parasite coevolution promotes innovation through deformations in fitness landscapes. eLife. 11:e76162.

25. Hancock RE, Reeves P. 1976. Lipopolysaccharide-deficient, bacteriophage-resistant mutants of *Escherichia coli* K-12. J Bacteriol. 127:98–108.

26. Hermisson J, Pennings PS. 2005. Soft sweeps: molecular population genetics of adaptation from standing genetic variation. Genetics. 169:2335–2352.

27. Inagaki M, Tanaka A, Suzuki R, Wakashima H, Kawaura T, Karita S, Nishikawa S, Kashimura N. 2000. Characterization of the binding of spike H protein of bacteriophage ΦX174 with receptor lipopolysaccharides. J Biochem. 127:577–583.

28. Incardona NL, Selvidge L. 1973. Mechanism of adsorption and eclipse of bacteriophage ΦX174. II. Attachment and eclipse with isolated Escherichia coli cell wall lipopolysaccharide. J Virol. 11:775–782.

29. Incardona NL, Tuech JK, Murti G. 1985. Irreversible binding of phage ΦX174 to cell-bound lipopolysaccharide receptors and release of virus-receptor complexes. Biochemistry. 24:6439– 6446.

30. Jamet A, Touchon M, Ribeiro-Gonçalves B, Carriço JA, Charbit A, Nassif X, Ramirez M, Rocha EPC. 2017. A widespread family of polymorphic toxins encoded by temperate phages. BMC Biol. 15:75.

31. Kawaura T, Inagaki M, Karita S, Kato M, Nishikawa S, Kashimura N. 2000. Recognition of receptor lipopolysaccharides by spike G protein of bacteriophage ΦX174. Biosci Biotechnol Biochem. 64:1993–1997.

32. Klein G, Müller-Loennies S, Lindner B, Kobylak N, Brade H, Raina S. 2013. Molecular and structural basis of inner core lipopolysaccharide alterations in *Escherichia coli*: incorporation of glucuronic acid and phosphoethanolamine in the heptose region. J Biol Chem. 288:8111–8127.

33. Klein G, Raina S. 2019. Regulated assembly of LPS, its structural alterations and cellular response to LPS defects. Int J Mol Sci. 20:356.

34. Kneidinger B, Marolda C, Graninger M, Zamyatina A, McArthur F, Kosma P, Valvano MA, Messner P. 2002. Biosynthesis pathway of ADP-L-glycero-β-D-manno-heptose in *Escherichia coli*. J Bacteriol. 184:7.

35. Kojima H, Inagaki M, Tomita T, Watanabe T. 2010. Diversity of non-stoichiometric substitutions on the lipopolysaccharide of *E. coli* C demonstrated by electrospray ionization single quadrupole mass spectrometry. Rapid Commun Mass Spectrom. 24:43–48.

36. Król JE, Hall DC, Balashov S, Pastor S, Sibert J, McCaffrey J, Lang S, Ehrlich RL, Earl J, Mell JC, et al. 2019. Genome rearrangements induce biofilm formation in *Escherichia coli* C – an old model organism with a new application in biofilm research. BMC Genomics. 20:767.

37. Krüger A, Lucchesi PMA. 2015. Shiga toxins and stx phages: highly diverse entities. Microbiol Read Engl. 161:451–462.

38. Kulikov EE, Golomidova AK, Prokhorov NS, Ivanov PA, Letarov AV. 2019. High-throughput LPS profiling as a tool for revealing of bacteriophage infection strategies. Sci Rep. 9:2958.

39. Labrie SJ, Samson JE, Moineau S. 2010. Bacteriophage resistance mechanisms. Nat Rev Microbiol. 8:317–327.

40. Latino L, Midoux C, Hauck Y, Vergnaud G, Pourcel C. 2016. Pseudolysogeny and sequential mutations build multiresistance to virulent bacteriophages in *Pseudomonas aeruginosa*. Microbiol Read Engl. 162:748–763.

41. van der Ley P, de Graaff P, Tommassen J. 1986. Shielding of *Escherichia coli* outer membrane proteins as receptors for bacteriophages and colicins by O-antigenic chains of lipopolysaccharide. J Bacteriol. 168:449–451.

42. Lind PA, Libby E, Herzog J, Rainey PB. 2019. Predicting mutational routes to new adaptive phenotypes. eLife. 8:e38822.

43. Luria SE, Delbrück M. 1943. Mutations of bacteria from virus sensitivity to virus resistance. Genetics. 28:491–511.

44. Maldonado RF, Sá-Correia I, Valvano MA. 2016. Lipopolysaccharide modification in Gram-negative bacteria during chronic infection. FEMS Microbiol Rev. 40:480–493.

45. Matsuura M. 2013. Structural modifications of bacterial lipopolysaccharide that facilitate Gram-Negative bacteria evasion of host innate immunity. Front Immunol. 4:109.

46. McArthur F, Andersson CE, Loutet S, Mowbray SL, Valvano MA. 2005. Functional analysis of the glycero-manno-heptose 7-phosphate kinase domain from the bifunctional HldE protein, which is involved in ADP-L-glycero-D-manno-heptose biosynthesis. J Bacteriol. 187:5292– 5300.

47. McKenna R, Xia D, Willingmann P, IIag LL, Krishnaswamy S, Rossmann MG, Olson NH, Baker TS, Incardona NL. 1992. Atomic structure of single-stranded DNA bacteriophage ΦX174 and its functional implications. Nature. 355:137–143.

48. Meyer JR, Dobias DT, Weitz JS, Barrick JE, Quick RT, Lenski RE. 2012. Repeatability and contingency in the evolution of a key innovation in phage Lambda. Science. 335:428–432.

49. Michel A, Clermont O, Denamur E, Tenaillon O. 2010. Bacteriophage ΦX174’s ecological niche and the flexibility of its *Escherichia coli* lipopolysaccharide receptor. Appl Environ Microbiol. 76:7310– 7313.

50. Monteiro R, Pires DP, Costa AR, Azeredo J. 2019. Phage therapy: going temperate? Trends Microbiol. 27:368–378.

51. Moxon ER, Rainey PB, Nowak MA, Lenski RE. 1994. Adaptive evolution of highly mutable loci in pathogenic bacteria. Curr Biol. 4:24–33.

52. Murray CJ, Ikuta KS, Sharara F, Swetschinski L, Aguilar GR, Gray A, Han C, Bisignano C, Rao P, Wool E, et al. 2022. Global burden of bacterial antimicrobial resistance in 2019: a systematic analysis. The Lancet. 399:629–655.

53. Mutalik VK, Adler BA, Rishi HS, Piya D, Zhong C, Koskella B, Kutter EM, Calendar R, Novichkov PS, Price MN, et al. 2020. High-throughput mapping of the phage resistance landscape in *E. coli*. PLOS Biol. 18:e3000877.

54. Nikaido H. 2003. Molecular basis of bacterial outer membrane permeability revisited. Microbiol Mol Biol Rev. 67:593–656.

55. Oechslin F. 2018. Resistance development to bacteriophages occurring during bacteriophage therapy. Viruses. 10:351.

56. Pagnout C, Sohm B, Razafitianamaharavo A, Caillet C, Offroy M, Leduc M, Gendre H, Jomini S, Beaussart A, Bauda P, et al. 2019. Pleiotropic effects of *rfa*-gene mutations on *Escherichia coli* envelope properties. Sci Rep. 9:9696.

57. Picelli S, Björklund ÅK, Reinius B, Sagasser S, Winberg G, Sandberg R. 2014. Tn5 transposase and tagmentation procedures for massively scaled sequencing projects. Genome Res. 24:2033–2040.

58. Qian J, Garrett TA, Raetz CRH. 2014. *In vitro* assembly of the outer core of the lipopolysaccharide from *Escherichia coli* K-12 and *Salmonella typhimurium*. Biochemistry. 53:1250–1262.

59. Raetz CRH, Whitfield C. 2002. Lipopolysaccharide endotoxins. Annu Rev Biochem. 71:635–700.

60. Ramirez J, Guarner F, Bustos Fernandez L, Maruy A, Sdepanian VL, Cohen H. 2020. Antibiotics as major disruptors of gut microbiota. Front Cell Infect Microbiol. 10:572912

61. Remold SK, Rambaut A, Turner PE. 2008. Evolutionary genomics of host adaptation in vesicular stomatitis virus. Mol Biol Evol. 25:1138–1147.

62. Rokyta DR, Burch CL, Caudle SB, Wichman HA. 2006. Horizontal gene transfer and the evolution of microvirid coliphage genomes. J Bacteriol. 188:1134–1142.

63. Rubinstein A, Mizrachi Y, Bernstein L, Shliozberg J, Golodner M, Liu GQ, Ochs HD. 2000. Progressive specific immune attrition after primary, secondary and tertiary immunizations with bacteriophage ΦX174 in asymptomatic HIV-1 infected patients. AIDS Lond Engl. 14:F55–62.

64. Sanger, Coulsox AR, Friedmax T, Barrell BG, Browns L, Fiddes JC, Hctchisos CA, Sloconbe PM, Switi M. 1978. The nucleotide sequence of bacteriophage ΦX174. J Mol Biol. 125:225–246.

65. Sant DG, Woods LC, Barr JJ, McDonald MJ. 2021. Host diversity slows bacteriophage adaptation by selecting generalists over specialists. Nat Ecol Evol. 5:350–359.

66. Schnaitman CA, Klena JD. 1993. Genetics of lipopolysaccharide biosynthesis in enteric bacteria. Microbiol Rev. 57:655–682.

67. Seregina TA, Petrushanko IY, Shakulov RS, Zaripov PI, Makarov AA, Mitkevich VA, Mironov AS. 2022. The inactivation of LPS biosynthesis genes in *E. coli* cells leads to oxidative stress. Cells. 11:2667.

68. Simpson BW, Trent MS. 2019. Pushing the envelope: LPS modifications and their consequences. Nat Rev Microbiol. 17:403–416.

69. Sinsheimer RL. 1959. A single-stranded deoxyribonucleic acid from bacteriophage ΦX174. J Mol Biol. 1:43–53.

70. Sun Y, Roznowski AP, Tokuda JM, Klose T, Mauney A, Pollack L, Fane BA, Rossmann MG. 2017. Structural changes of tailless bacteriophage ΦX174 during penetration of bacterial cell walls. Proc Natl Acad Sci. 114:13708–13713.

71. Valvano MA, Marolda CL, Bittner M, Glaskin-Clay M, Simon TL, Klena JD. 2000. The *rfaE* gene from *Escherichia coli* encodes a bifunctional protein involved in biosynthesis of the lipopolysaccharide core precursor ADP-L-glycero-D-manno-heptose. J Bacteriol. 182:488–497.

72. Weisbeek PJ, Van de Pol JH, Van Arkel GA. 1973. Mapping of host range mutants of bacteriophage ΦX174. Virology. 52:408–416.

73. Whitfield C, Heinrichs DE, Yethon JA, Amor KL, Monteiro MA, Perry MB. 1999. Assembly of the R1-type core oligosaccharide of Escherichia coli lipopolysaccharide. 5:151–156.

74. WHO. 2021. Antimicrobial resistance. Available from: https://www.who.int/news-room/fact-sheets/detail/antimicrobial-resistance

75. Wichman HA, Badgett MR, Scott LA, Boulianne CM, Bull JJ. 1999. Different trajectories of parallel evolution during viral adaptation. Science .285:422–424.

76. Yethon JA, Heinrichs DE, Monteiro MA, Perry MB, Whitfield C. 1998. Involvement of *waaY*, *waaQ*, and *waaP* in the modification of *Escherichia coli* lipopolysaccharide and their role in the formation of a stable outer membrane. J Biol Chem. 273:26310–26316.

77. Yethon JA, Vinogradov E, Perry MB, Whitfield C. 2000. Mutation of the lipopolysaccharide core glycosyltransferase encoded by *waaG* destabilizes the outer membrane of *Escherichia coli* by interfering with core phosphorylation. J Bacteriol. 182:5620–5623.

78. Zhang G, Meredith TC, Kahne D. 2013. On the essentiality of lipopolysaccharide to Gram-negative bacteria. Curr Opin Microbiol. 16:779–785.

## References

79. Amor K, Heinrichs DE, Frirdich E, Ziebell K, Johnson RP, Whitfield C. 2000. Distribution of core oligosaccharide types in lipopolysaccharides from *Escherichia coli*. Infect Immun. 68:1116–1124.

80. Belunis CJ, Clementz T, Carty SM, Raetz CRH. 1995. Inhibition of lipopolysaccharide biosynthesis and cell growth following inactivation of the *kdtA* gene in *Escherichia coli*. J Biol Chem. 270:27646–27652.

81. Bohm K, Porwollik S, Chu W, Dover JA, Gilcrease EB, Casjens SR, McClelland M, Parent KN. 2018. Genes affecting progression of bacteriophage P22 infection in *Salmonella* identified by transposon and single gene deletion screens: host genes affecting phage P22 infection. Mol Microbiol. 108:288–305.

82. Bono LM, Gensel CL, Pfennig DW, Burch CL. 2012. Competition and the origins of novelty: experimental evolution of niche-width expansion in a virus. Biol Lett. 9:20120616– 20120616.

83. van Charante F, Holtappels D, Blasdel B, Burrowes B. 2019. Isolation of bacteriophages. In: Harper DR, Abedon ST, Burrowes BH, McConville ML, editors. Bacteriophages: Biology, Technology, Therapy. Springer, Cham. p. 1–32.

84. Fang J, Wei Y. 2011. Expression, purification and characterization of the *Escherichia coli* integral membrane protein YajC. Protein Pept Lett. 18:601–608.

85. Frey PA. 1996. The Leloir pathway: a mechanistic imperative for three enzymes to change the stereochemical configuration of a single carbon in galactose. FASEB j. 10:461–470.

86. Genevaux P, Bauda P, DuBow MS, Oudega B. 1999. Identification of Tn10 insertions in the rfaG, rfaP, and galU genes involved in lipopolysaccharide core biosynthesis that affect Escherichia coli adhesion. Arch of Microbiol. 172:1–8.

87. Hancock RE, Reeves P. 1976. Lipopolysaccharide-deficient, bacteriophage-resistant mutants of *Escherichia coli* K-12. J Bacteriol. 127:98–108.

88. Heinrichs DE, Yethon JA, Amor PA, Whitfield C. 1998. The assembly system for the outer core portion of R1- and R4-type lipopolysaccharides of *Escherichia coli*: the R1 core-specific β-glucosyltransferase provides a novel attachment site for O-polysaccharides. J Biol Chem. 273:29497–29505.

89. Heinrichs DE, Yethon JA, Whitfield C. 1998. Molecular basis for structural diversity in the core regions of the lipopolysaccharides of *Escherichia coli* and *Salmonella enterica*. Mol Microbiol. 30:221–232.

90. Jansson P-E, Lindberg B, Lindberg AA, Wollin R. 1981. Structural studies on the hexose region of the core in lipopolysaccharides from Enterobacteriaceae. Eur J Biochem. 115:571–577.

91. Karp PD, Billington R, Caspi R, Fulcher CA, Latendresse M, Kothari A, Keseler IM, Krummenacker M, Midford PE, Ong Q, et al. 2019. The BioCyc collection of microbial genomes and metabolic pathways. Brief Bioinform. 20:1085–1093.

92. Kawaura T, Inagaki M, Karita S, Kato M, Nishikawa S, Kashimura N. 2000. Recognition of receptor lipopolysaccharides by spike G protein of bacteriophage ΦX174. Biosci Biotechnol Biochem. 64:1993–1997.

93. Klein G, Lindner B, Brabetz W, Brade H, Raina S. 2009. *Escherichia coli* K-12 suppressor-free mutants lacking early glycosyltransferases and late acyltransferases: minimal lipopolysaccharide structure and induction of envelope stress response. J Biol Chem. 284:15369–15389.

94. Klein G, Müller-Loennies S, Lindner B, Kobylak N, Brade H, Raina S. 2013. Molecular and structural basis of inner core lipopolysaccharide alterations in *Escherichia coli*: incorporation of glucuronic acid and phosphoethanolamine in the heptose region. J Biol Chem. 288:8111– 8127.

95. Kneidinger B, Marolda C, Graninger M, Zamyatina A, McArthur F, Kosma P, Valvano MA, Messner P. 2002. Biosynthesis pathway of ADP-L-glycero-β-D-manno-heptose in *Escherichia coli*. J. Bacteriol. 184:363–369.

96. Król JE, Hall DC, Balashov S, Pastor S, Sibert J, McCaffrey J, Lang S, Ehrlich RL, Earl J, Mell JC, et al. 2019. Genome rearrangements induce biofilm formation in *Escherichia coli* C – an old model organism with a new application in biofilm research. BMC Genomics 20:767.

97. Kulikov EE, Golomidova AK, Prokhorov NS, Ivanov PA, Letarov AV. 2019. High-throughput LPS profiling as a tool for revealing of bacteriophage infection strategies. Sci Rep. 9:2958.

98. Labrie SJ, Samson JE, Moineau S. 2010. Bacteriophage resistance mechanisms. Nat Rev Microbiol. 8:317–327.

99. Leipold MD, Vinogradov E, Whitfield C. 2007. Glycosyltransferases involved in biosynthesis of the outer core region of *Escherichia coli* lipopolysaccharides exhibit broader substrate specificities than is predicted from lipopolysaccharide structures. J Biol Chem. 282:26786– 26792.

100. van der Ley P, de Graaff P, Tommassen J. 1986. Shielding of *Escherichia coli* outer membrane proteins as receptors for bacteriophages and colicins by O-antigenic chains of lipopolysaccharide. J Bacteriol 168:449–451.

101. Matsuura M. 2013. Structural modifications of bacterial lipopolysaccharide that facilitate Gram-Negative bacteria evasion of host innate immunity. Front Immunol. 109:4.

102. McArthur F, Andersson CE, Loutet S, Mowbray SL, Valvano MA. 2005. Functional analysis of the glycero-manno-heptose 7-phosphate kinase domain from the bifunctional HldE protein, which is involved in ADP-L-glycero-D-manno-heptose biosynthesis. J Bacteriol. 187:5292–5300.

103. Mutalik VK, Adler BA, Rishi HS, Piya D, Zhong C, Koskella B, Kutter EM, Calendar R, Novichkov PS, Price MN, et al. 2020. High-throughput mapping of the phage resistance landscape in *E. coli*. PLoS Biol. 18:e3000877.

104. Pagnout C, Sohm B, Razafitianamaharavo A, Caillet C, Offroy M, Leduc M, Gendre H, Jomini S, Beaussart A, Bauda P, et al. 2019. Pleiotropic effects of *rfa*-gene mutations on *Escherichia coli* envelope properties. Sci Rep. 9:9696.

105. Pierson DE, Carlson S. 1996. Identification of the *galE* gene and a *galE* homolog and characterization of their roles in the biosynthesis of lipopolysaccharide in a serotype O:8 strain of *Yersinia enterocolitica*. J Bacteriol. 178:5916–5924.

106. Raetz CRH, Whitfield C. 2002. Lipopolysaccharide endotoxins. Annu Rev Biochem. 71:635–700.

107. Schnaitman CA, Austin EA. 1990. Efficient incorporation of galactose into lipopolysaccharide by *Escherichia coli* K-12 strains with polar *galE* mutations. J Bacteriol. 172:5511–5513.

108. Schnaitman CA, Klena JD. 1993. Genetics of lipopolysaccharide biosynthesis in enteric bacteria. Microbiol Rev. 57:655–682.

109. Schulze RJ, Komar J, Botte M, Allen WJ, Whitehouse S, Gold VAM, Nijeholt JAL a, Huard K, Berger I, Schaffitzel C, et al. 2014. Membrane protein insertion and proton-motive-force-dependent secretion through the bacterial holo-translocon SecYEG–SecDF–YajC–YidC. PNAS 111:4844–4849.

110. Vinogradov EV, van der Drift K, Thomas-Oates JE, Meshkov S, Brade H, Holst O. 1999. The structures of the carbohydrate backbones of the lipopolysaccharides from *Escherichia coli* rough mutants F470 (R1 core type) and F576 (R2 core type): LPS from *E. coli* R1 and R2 core types. Eur J Biochem. 261:629–639.

111. Weissborn AC, Liu Q, Rumley MK, Kennedy EP. 1994. UTP:α-D-glucose-L-phosphate uridylyltransferase of *Escherichia coli*: isolation and DNA Sequence of the *galU* gene and purification of the enzyme. J Bacteriol. 176:2611–2618.

112. Whitfield C, Heinrichs DE, Yethon JA, Amor KL, Monteiro MA, Perry MB. 1999. Assembly of the R1-type core oligosaccharide of *Escherichia coli* lipopolysaccharide. J Endotoxin Res. 5:151–156.

113. Yethon JA, Heinrichs DE, Monteiro MA, Perry MB, Whitfield C. 1998. Involvement of *waaY*, *waaQ*, and *waaP* in the modification of *Escherichia coli* lipopolysaccharide and their role in the formation of a stable outer membrane. J Biol Chem. 273:26310–26316.

114. Yethon JA, Vinogradov E, Perry MB, Whitfield C. 2000. Mutation of the lipopolysaccharide core glycosyltransferase encoded by *waaG* destabilizes the outer membrane of *Escherichia coli* by interfering with core phosphorylation. J Bacteriol. 182:5620–5623.

